# Rest promotes microglial clearance of degenerating neurons

**DOI:** 10.64898/2026.11.16.766874

**Authors:** Zongqin Xiang, Bin Mou, Miaoting Li, Junliang Chen, Lang Huang, Yong U. Liu

## Abstract

How the brain coordinates the removal of degenerating neurons with host rest and activity remains poorly understood. Using longitudinal two-photon imaging in mice with cytosolic TDP-43 proteinopathy, we show that microglia remove neuronal corpses through an ordered sequence of reaching, approaching, isolating and digesting. Clearance was higher during the light (resting) phase and lower during the dark (active) phase. This pattern reversed after light–dark cycle inversion and was blunted by sleep disruption. Chemogenetic inhibition of locus coeruleus noradrenergic neurons increased engulfment during the active phase, while activation and inhibition of the β2-adrenergic receptor ADRB2 suppressed and enhanced clearance, respectively, identifying noradrenergic signaling as a brake on engulfment. However, sustained microglial ADRB2 deletion impaired clearance and blunted light–dark differences. These findings reveal that microglial removal of degenerating neurons is enhanced during rest and support a role for noradrenergic signaling in coordinating this cellular maintenance process with the host’s rest–activity cycle.

## Introduction

The removal of dying cells is essential for tissue homeostasis. Across organs, neighboring cells and professional phagocytes recognize and eliminate cellular corpses through tightly regulated mechanisms^1,2^. Failure of clearance can promote persistent inflammation and tissue dysfunction^3,4^. In the central nervous system (CNS), this task presents distinctive challenges: most neurons have limited capacity for replacement and are embedded within specialized networks that support information processing, behavior and memory^5,6^. Removing degenerating neurons therefore requires spatially and temporally coordinated cellular responses. How this local maintenance process is coordinated with changes in the host’s rest– activity state remains poorly understood.

Microglia, the resident macrophages of the CNS, continuously survey the brain parenchyma and remove dying cells and damaged cellular components^7–11^. The importance of glial corpse clearance is further supported by studies in Drosophila, in which disruption of glial phagocytic machinery leads to the accumulation of apoptotic neurons and progressive neurodegeneration^12^. Studies of macrophage and microglial phagocytosis have defined key molecular steps, including responses to ‘find-me’ signals, recognition of ‘eat-me’ signals and phagolysosomal degradation^13–15^. However, fixed-tissue and culture-based approaches cannot fully resolve how these events unfold as individual neurons degenerate in the intact brain. The sequence of microglial engagement, its relationship to transcriptional state and the factors governing progression from initial contact to completed corpse removal remain incompletely understood.

Beyond local signals from dying neurons, microglial clearance may be influenced by the physiological state of the host. Engulfment and degradation of an entire neuronal soma involve extensive cytoskeletal remodeling, membrane trafficking and lysosomal processing, raising the question of whether this cellular activity varies between rest and activity. The sleep–wake cycle is accompanied by changes in neural activity and neuromodulatory tone. In particular, norepinephrine (NE), a major regulator of arousal and vigilance, is generally higher during wakefulness and lower during sleep^16,17^. In vivo imaging studies have shown that microglial process dynamics vary across wakefulness, sleep and anesthesia^18–20^, providing a basis for investigating whether arousal-related signals also regulate neuronal corpse removal. Whether such regulation extends from microglial surveillance to the sustained process of engulfing and clearing degenerating neurons remains unknown.

Here we used a mouse model of cytosolic TDP-43 proteinopathy, a pathological feature implicated in several neurodegenerative disorders^21–25^, to track neuronal degeneration and microglial clearance longitudinally. Two-photon imaging in awake mice resolved an ordered sequence of reaching, approaching, isolating and digesting (RAID), while cellular behavior-guided transcriptomics associated microglial engagement categories with distinct transcriptional profiles. Clearance was higher during the light (resting) phase, followed the shifted light–dark schedule after cycle inversion and showed reduced phase dependence following sleep disruption. Resting-phase clearance was accompanied by greater enrichment of lysosomal and metabolic gene programs. Noradrenergic interventions identified NE– ADRB2 signaling as a brake on engulfment. Sustained microglial ADRB2 deletion nevertheless impaired clearance and blunted light–dark differences, distinguishing the consequences of receptor loss from the effects of pharmacological inhibition. Together, these findings link microglial removal of degenerating neurons to the host’s rest–activity cycle and identify noradrenergic signaling as a regulator of this cellular maintenance process.

## Results

### Longitudinal in vivo imaging captures microglial phagocytosis of degenerating neurons

To investigate microglial responses to progressive neurodegeneration in vivo, we expressed a nuclear localization sequence-mutant TDP-43 fused to mScarlet (TDP-43ΔNLS-mScarlet) in microglial reporter mice using an adeno-associated virus (AAV), with mScarlet alone as a control^26,27^ **(Figure 1A)**. Longitudinal two-photon imaging in awake, head-fixed mice on a treadmill enabled repeated tracking of neurons and neighboring microglia across multiple timescales **(Figure 1B; Figure S1A)**. Histological analysis showed that TDP-43ΔNLS-mScarlet expression induced cytosolic TDP-43 proteinopathy (cytoTDP-43) in the primary motor cortex, with TDP-43 accumulation and a progressive reduction in NeuN-positive neurons **(Figure S2A-D)**. By 21 days after injection, neuronal loss was accompanied by microglial proliferation and increased CD68 expression; these changes were absent in controls **(Figure S2E and S2F)**. Complementary in vivo calcium imaging showed that reduced neuronal calcium-event frequency and amplitude preceded cell loss **(Figure S3)**.

**Figure 1.**
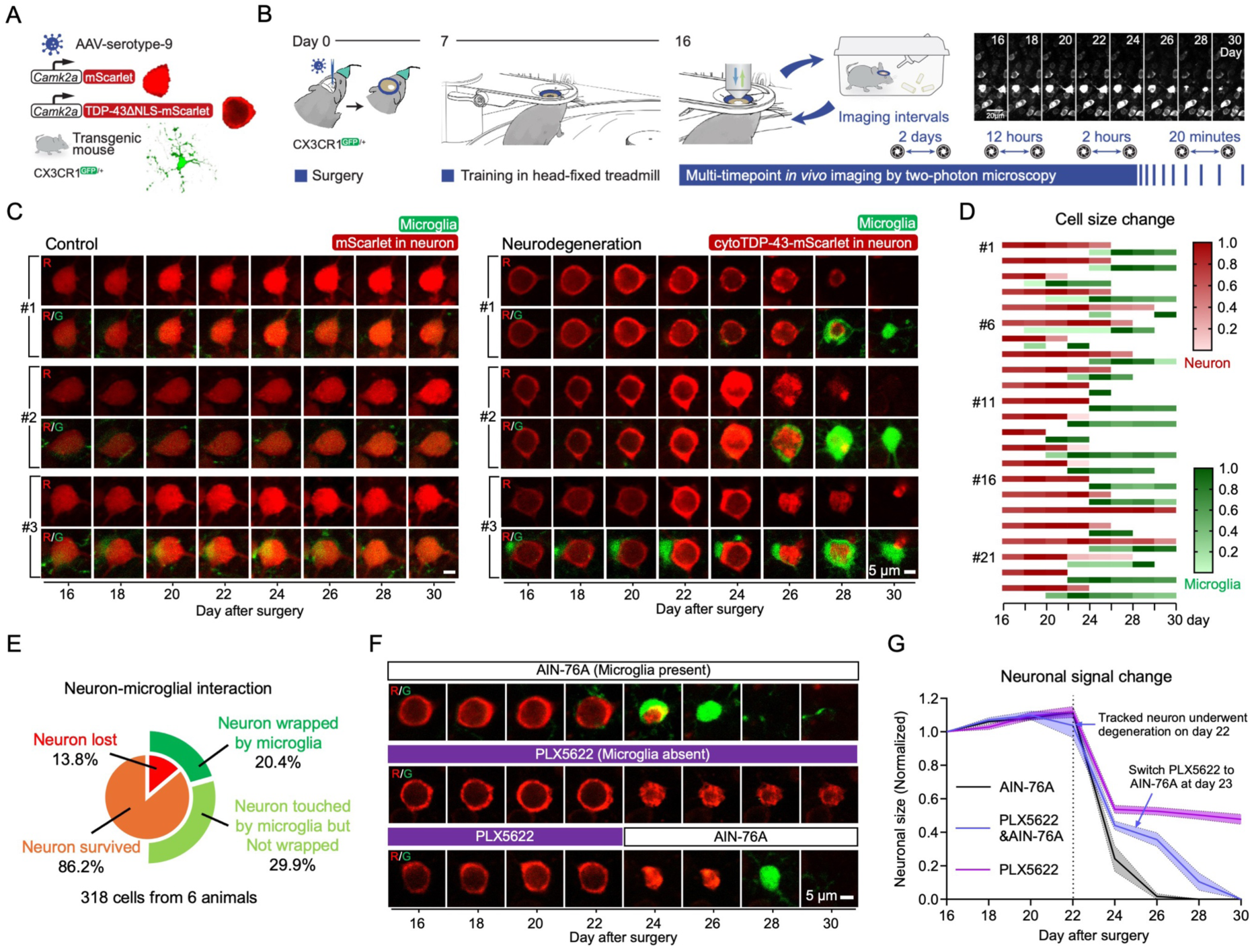
Longitudinal in vivo two-photon imaging reveals progressive neurodegeneration and microglial responses in cytoTDP-43-overexpressing cortex. **(A)** Experimental paradigm combining AAV-mediated gene delivery and transgenic reporter mice for longitudinal two-photon imaging. AAV vectors under the *Camk2α* promoter expressed either mScarlet (Control, Ctrl) or TDP43-mtNLS-linker-mScarlet (Neurodegeneration, ND) selectively in excitatory neurons. **(B)** Workflow depicting chronic cranial window implantation, stereotaxic AAV injection and longitudinal imaging. Mice were habituated to a head-fixed treadmill to minimize motion artifacts. **(C)** Repeated two-photon imaging at 2-day intervals of individual neurons from 16-30 days after surgery. In controls (left), mScarlet signal intensified without morphological alteration. In cytoTDP-43 neurons (right), TDP43-mScarlet initially localized to the cytoplasm, subsequently translocated into the nucleus preceding cell death and accompanied by microglial engulfment. Scale bar, 5 μm. **(D)** Temporal profiles of 23 neuronal degeneration events, aligned to microglial recruitment and engagement across 5 mice. **(E)** Pie charts summarizing survival outcomes: all lost neurons were phagocytized by microglia; a subset of microglia-neuron interactions resolved without neuronal death. **(F)** Imaging under pharmacological microglial depletion (PLX5622): neuronal corpses persisted in the absence of microglia (middle) but were swiftly eliminated upon microglial repopulation following PLX5622 withdrawal (bottom). Scale bar, 5 μm. **(G)** Morphometric analysis of neuronal soma size change during degeneration (selected neurons initiation degeneration at day 22) with or without microglial surveillance. Trend lines represent the mean ± s.e.m. n=14 neurons from 4 animals for AIN-76A group, n=10 neurons from 3 animals for PLX5622 and PLX5622&AIN-76A group. Detailed statistics are provided in Supplementary Table1.

We began longitudinal imaging 16 days after surgery, allowing recovery from viral injection and cranial-window implantation, and tracked the same neurons and microglia every two days. Control neurons and microglial organization remained stable, whereas cytoTDP-43-expressing neurons exhibited progressive protein accumulation, somatic shrinkage and eventual loss of fluorescence associated with microglial engulfment **(Figure S1B; Movie 1)**. Declining neuronal survival was accompanied by transient local increases in microglial density **(Figure S1C)**. High-resolution tracking of individual neurons showed progressive cytoplasmic cytoTDP-43 accumulation followed by microglial recruitment and engulfment, whereas control neurons maintained a uniform distribution of mScarlet fluorescence **(Figure 1C)**. Alignment of 23 neuronal degeneration events revealed coordinated changes in neuronal morphology and microglial engagement: as neuronal somata shrank, microglia accumulated around the targets and enlarged during engulfment **(Figure 1D)**. Many transient microglia– neuron contacts resolved without neuronal loss, indicating that contact alone did not predict terminal degeneration **(Figure 1E)**.

To test whether microglia were required for corpse removal, we depleted them pharmacologically with PLX5622^28^. Neuronal somata continued to shrink and collapse, but cellular remnants persisted in the absence of microglia. Following drug withdrawal, repopulating microglia rapidly removed these remnants **(Figure. 1F and 1G; Figure S4A-C; Movie 2)**. Microglial depletion was also associated with greater neuronal loss in the cytoTDP-43 cortex **(Figure S4D and S4E)**. These experiments distinguish neuronal structural collapse from subsequent corpse removal and establish a requirement for microglia in the latter process.

### Cellular behavior-guided transcriptomics identifies molecular programs associated with microglial engulfment

To resolve the sequence of microglial clearance, we performed time-lapse imaging at intervals of two hours, 20 minutes and three minutes **(Movie 3)**. This revealed four ordered behaviors: reach, involving process extension towards the target; approach, involving somatic migration; isolate, involving complete encapsulation; and digest, involving degradation or incorporation of neuronal material. We termed this sequence RAID **(Figure 2A)**. Transmission electron microscopy identified ultrastructural features consistent with approach, isolation and digestion **(Figure 2B)**.

**Figure 2.**
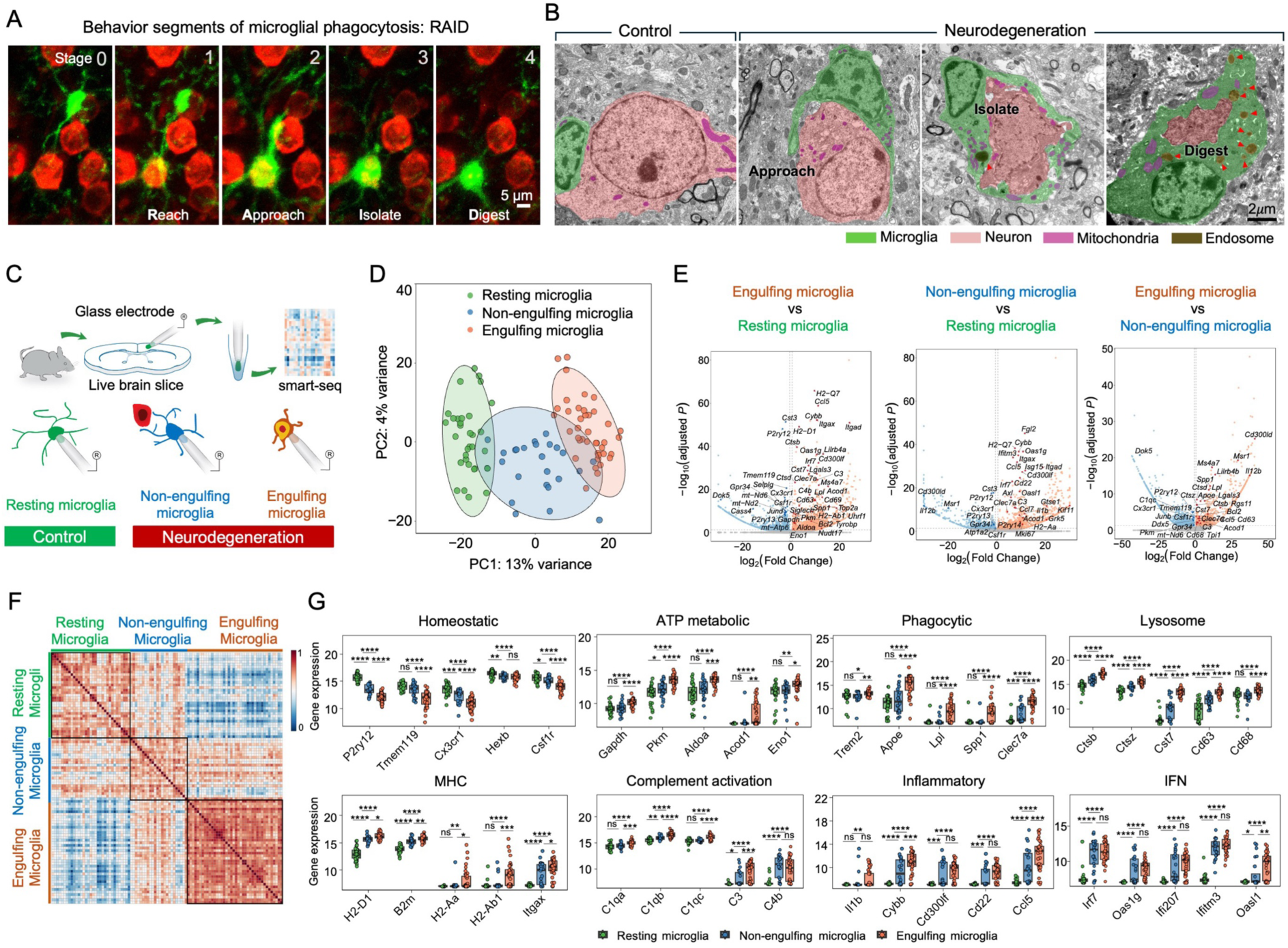
Cellular behavior-guided (CBG) Smart-seq reveals stage-associated transcriptional states during microglial neuronal clearance. **(A)** Schematic illustrating five sequential stages of microglial engagement with degenerating neurons defined by longitudinal in vivo imaging: Stage 0 (Prepare), initial contact; Stage 1 (Reach), extension of microglial processes toward the neuronal soma; Stage 2 (Approach), migration of the microglial soma; Stage 3 (Isolate), complete encapsulation of the neuronal soma; and Stage 4 (Digest), degradation and clearance of neuronal debris. Scale bars, 5 μm. **(B)** Representative transmission electron microscopy images obtained three weeks after surgery, validating ultrastructural features corresponding to microglial approach, isolation, and digestion of neuronal remnants. Scale bars, 2 μm. **(C)** Imaging-guided smart-seq workflow. Based on real-time morphology and engagement with degenerating neurons, patched microglia were classified as resting microglia from control cortex, non-engulfing microglia and engulfed microglial from TDP-43 cortex. **(D)** Principal component analysis (PCA) shows separation of resting microglia (green), non-engulfing microglia (blue), and engulfing microglia (orange) by smart-seq, each dot represents one cell. **(E)** Volcano plots showing the differentially expressed genes (DEGs) between the indicated microglia states in (D). Benjamini-Hochberg (BH) test was used for statistical testing. **(F)** Correlation heatmap based on the top 100 selected genes, showing transcriptomic differences among resting-state, non-engulfing and engulfing microglia. **(G)** Expression of the indicated marker genes across the three microglial states. Individual cells are shown. Statistical significance was assessed by one-way ANOVA followed by Tukey’s HSD post hoc test. Detailed statistics are provided in Supplementary Table1.

To characterize the molecular programs associated with microglial engagement, we performed cellular behavior-guided single-cell transcriptomics (CBG-seq). Individual microglia were isolated from acute brain slices according to microscopically defined engagement status— resting, activated but non-engulfing, or engulfing—and subjected to Smart-seq profiling **(Figure 2C)**. Here, “resting” denotes the cellular engagement category rather than the animal’s behavioral state. The engulfing category included cells involved in target isolation and digestion; thus, the sampling distinguished engagement categories rather than each RAID stage individually. Principal component analysis placed non-engulfing microglia between resting and engulfing cells, while gene-expression correlations supported clustering by engagement status **(Figure 2D and 2F)**.

Engulfing microglia showed greater expression of genes associated with phagocytosis, lysosomal function, antigen presentation, ATP metabolism and glycolysis, together with reduced homeostatic signatures **(Figure 2E and 2G; Figure S5A)**. Weighted gene co-expression network analysis of the 5,000 most highly expressed genes identified an engulfment-associated module containing phagocytic genes (*Tyrobp*, *Lgals3*), complement genes (*C1qa*, *C1qb*), lysosomal genes (*Ctsb*, *Ctsd*, *Cd63*) and glycolytic genes (*Aldoa*, *Eno1*). A module containing the homeostatic genes *P2ry12*, *Cx3cr1* and *Csf1r* showed reduced expression in engulfing cells **(Figure S5B and S5C)**. Together, these findings link behavior-defined engagement states to distinct transcriptional profiles, with engulfing microglia showing greater expression of phagocytic, lysosomal and metabolic gene programs and reduced homeostatic signatures.

### Microglial clearance is enhanced during the resting phase

Having resolved the behavioral sequence of neuronal clearance, we asked whether its progression varied across the host’s rest–activity cycle. Imaging at two-hour intervals across a single night revealed temporal fluctuations in microglial phagocytic activity **(Figure S6A)**. To limit perturbation from repeated imaging, we compared clearance at representative times in the light (resting) and dark (active) phases, at 08:00 and 20:00, respectively **(Figure 3A)**. Under the normal light–dark schedule, microglia engulfed more degenerating neurons, completed clearance more efficiently and progressed further through RAID during the resting phase. Engulfment and clearance completion were reduced during the active phase **(Figure 3B-D; Figure S6B; Movie 4)**.

**Figure 3.**
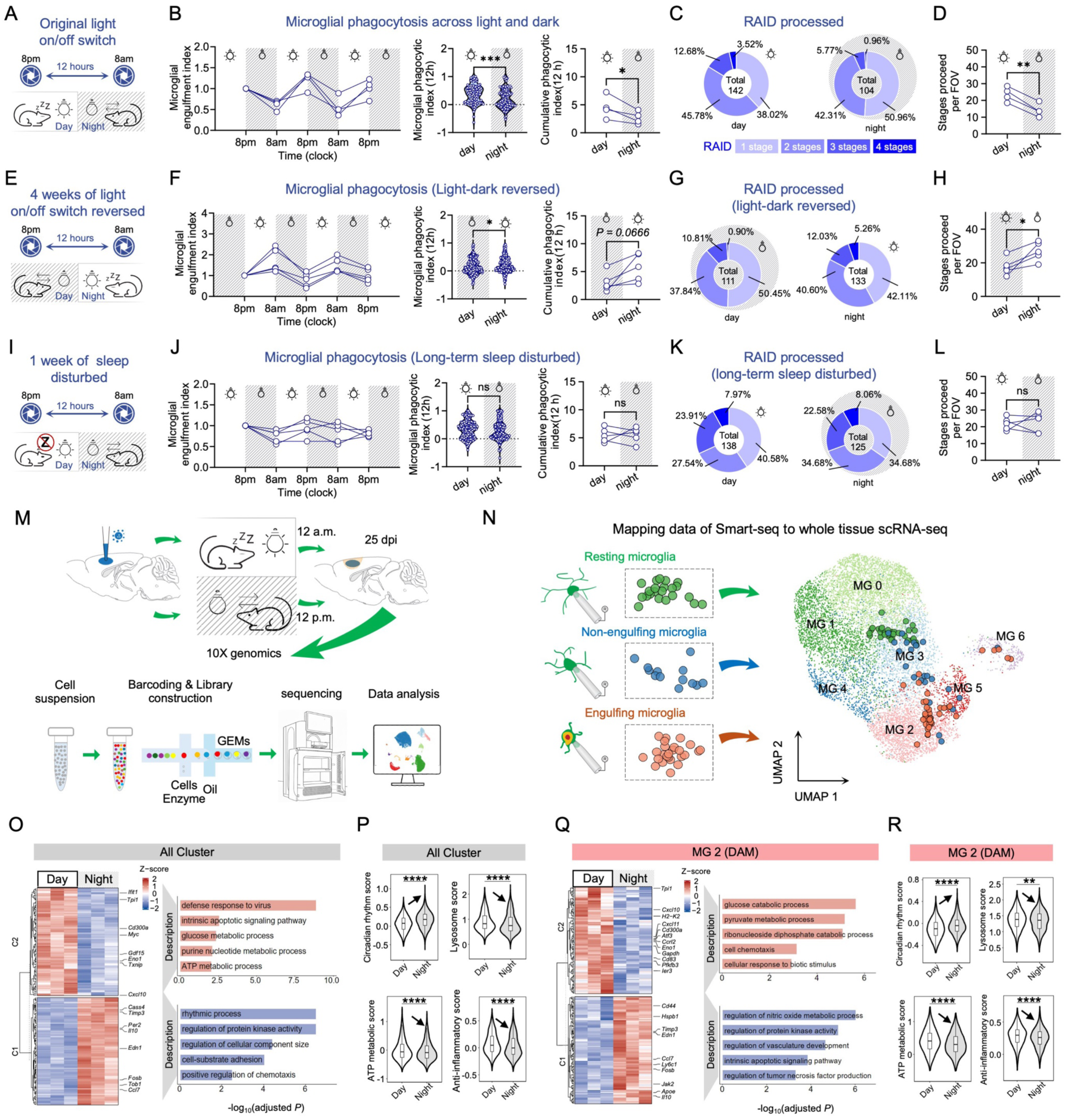
Circadian regulation of microglial phagocytic activity during neuronal degeneration. **(A, E, I)** Experimental design of in vivo two-photon imaging under three conditions: normal circadian cycle (A), reversed circadian cycle (E), and sleep disturbance (I). Imaging sessions were conducted at 8:00 a.m. and 8:00 p.m. **(B)** Under a normal light-dark cycle, the number of engulfed neurons (B, left), microglial phagocytic index (B, middle; 1 indicates successful clearance of one neuron), and cumulative phagocytic index per field of view (FOV; B, right) were significantly higher during the day and reduced at night. **(C, D)** During the daytime, microglia more frequently progressed through all four stages of engulfment (c) and advanced more stages (d). **(F-L)** These day-night differences were inverted under the reversed circadian cycle (F-H) and markedly attenuated under sleep disturbance (J-L). n=4 animals for normal circadian group, n=5 animals for other two groups; For the middle panels in (B, F, J), n represents individual event pooled from 4 or 5 animals. **(M)** Experimental design for single-cell RNA sequencing of cortical cells collected at 12:00 (day) and 24:00 (night). **(N)** UMAP visualization of microglial subclusters (MG0–MG6) in the independently generated rest–active scRNA-seq atlas, with CBG-seq-defined resting-state, non-engulfing and engulfing profiles projected onto the reference. Actively engulfing microglia preferentially mapped to the DAM-like/engulfment-associated MG2 population in this dataset. **(O-R)** Heatmap visualization of day–night DEGs and GO biological process enrichment analysis of significantly upregulated genes and downregulated genes in total microglia (O) and DAM-like/engulfment-associated MG2 population (Q); Violin plots showing signature scores of Circadian rhythms, ATP metabolic, Lysosome and Anit-inflammatory scores among day and night in total microglia (P) and DAM-like/engulfment-associated MG2 population (DAM; MG2; R). *, *P*<0.05, **, *P*<0.01, ****, *P*<0.0001; unpaired *t*-test (two-tailed). Detailed statistics are provided in Supplementary Table1.

After four weeks under an inverted light–dark schedule, the clearance pattern reversed relative to clock time and followed the shifted resting and active phases **(Figure 3E-H; Figure S6C)**. Sleep disruption during the resting phase markedly attenuated the light–dark difference in clearance **(Figure 3I-L)**. These experiments identify the resting phase as a period of enhanced neuronal clearance and show that the phase-dependent pattern is sensitive to both schedule inversion and sleep disruption.

We next examined whether these functional differences were accompanied by changes in microglial transcriptional profiles. Cortical tissue was collected during the light/resting phase at 12:00 and the dark/active phase at 24:00 for single-cell RNA sequencing **(Figure 3M)**. The dataset comprised 71,269 cells assigned to twelve major cell types; microglia and endothelial cells accounted for 31.3% and 40.8% of captured cells, respectively **(Figure S7A and S7B)**. Microglial subclustering identified seven populations (MG0–MG6), annotated as homeostatic, inflammatory-primed, disease-associated, metabolically active, pro-inflammatory, interferon-responsive and proliferative, respectively **(Figure S7C and S7E)**. Pseudotime, RNA velocity and module-scoring analyses suggested relationships among these populations and identified differences in phagocytic, lysosomal, metabolic and immune-associated gene programs **(Figure S7D and S7F-H)**. Integration with CBG-seq profiles associated the three engagement categories with distinct regions of the transcriptional landscape. The engulfing signature mapped predominantly to the disease-associated microglia (DAM)-like MG2 population **(Figure 3N)**.

Differential-expression and pathway analyses identified phase-associated differences in cellular defense, apoptotic signaling, metabolism, kinase signaling, adhesion and chemotactic programs **(Figure 3O)**. Signature scoring showed greater enrichment of ATP metabolism-associated, lysosomal and anti-inflammatory programs during the resting phase, with stronger rhythmicity-associated signatures during the active phase **(Figure 3P)**. Within MG2, which was enriched for the CBG-seq engulfment signature, metabolic and lysosomal gene programs were stronger during the resting phase, whereas inflammatory signatures were enhanced during the active phase **(Figure 3Q and 3R)**.

The relative representation of microglial populations also differed between phases. Homeostatic MG0 and metabolically active MG3 were enriched during the resting phase, whereas inflammatory-primed MG1 and pro-inflammatory MG4 were enriched during the active phase **(Figure S8A-C)**. MG1 expressed higher levels of inflammatory genes, including *Marcksl1, Nlrp3, Il1b, Cd14, Nfkb1 and Ccl7*, while MG0 preferentially expressed genes associated with metabolic maintenance and homeostasis **(Figure S8D and S8E)**. MG0 also exhibited stronger phagocytic and lysosomal signatures and weaker inflammatory signatures than MG1 **(Figure S8F and S8G)**. Thus, the active phase was associated with greater representation of populations bearing inflammatory and stress-responsive signatures ^29,30^, whereas the resting phase was associated with populations showing stronger homeostatic and metabolic signatures. Together, these analyses show that enhanced resting-phase clearance is accompanied by differences in both microglial population composition and lysosomal and metabolic gene programs. We next investigated whether arousal-related signals contribute to the regulation of clearance observed in vivo.

### Noradrenergic signaling restrains microglial clearance during the host active phase

We focused on NE as a candidate link between arousal and microglial clearance. Our previous work showed that reduced NE during anesthesia accompanies increased microglial process dynamics, whereas elevated NE suppresses these responses^18^. To determine whether this regulation extends to neuronal engulfment, we imaged microglia during reversible transitions between wakefulness and general anesthesia in the host active phase **(Figure 4A)**. An in vivo NE sensor showed a rapid decline in signal following anesthesia induction and a rebound during emergence **(Figure 4B; Figure S9A)**. Microglial phagocytic activity increased after anesthesia onset and declined following recovery of wakefulness **(Figure 4C and 4D; Figure S9B; Movie 5)**.

**Figure 4.**
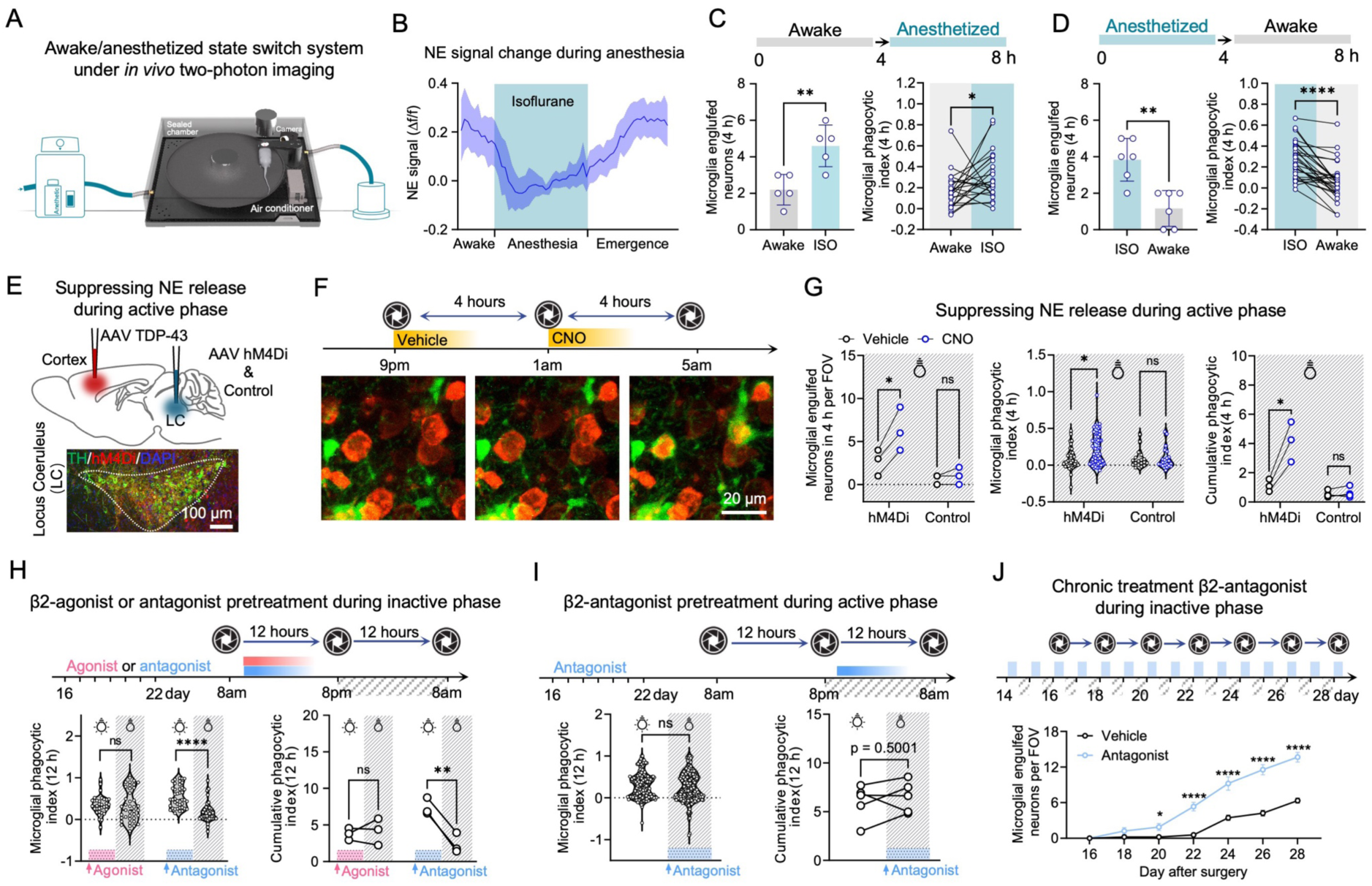
NE-Adrb2 signaling regulates daily fluctuations in microglial phagocytosis of neuronal corpses. **(A)** Experimental design for in vivo two-photon imaging of microglial phagocytosis under awake versus anesthetized conditions. **(B)** Time course of NE sensor signals during transitions from wakefulness to anesthesia and subsequent emergence. Trend lines represent the mean ± s.e.m. n= 3 animals. **(C-D)** Quantification of microglial phagocytic activity under awake versus anesthetized conditions, showing enhanced engulfment during anesthesia and suppression upon emergence. n=5 animals for (C) left panel, n=6 animals for (D)left panel**, n**=24 cells from 5 animals for (C) right panel, n=30 cells from 6 animals for (D) right panel. **(E)** Experimental design for chemogenetic inhibition of locus coeruleus (LC) noradrenergic neurons. Scale bars, 100 μm. **(F)** Representative images showing microglial engulfment of degenerating neurons following CNO administration. Scale bars, 20 μm. **(G)** Quantification of microglial phagocytosis before and after CNO administration, showing no effect in control mice but a significant increase following LC inhibition. n=3 animals for each group, n represents individual event pooled from 3 animals for middle panel. **(H)** Pharmacological manipulation of Adrb2 at 10:00 a.m. Activation of Adrb2 attenuated circadian differences in microglial phagocytosis, whereas Adrb2 inhibition enhanced day-night differences. n=3 animals for right panel, n represents individual event pooled from 3 animals for left panel. **(I)** Adrb2 antagonist administration at 10:00 p.m. attenuated day-night differences in microglial phagocytic activity. n=5 animals for right panel, n represents individual event pooled from 5 animals for left panel. **(J)** Long-term daily treatment with a β2-adrenergic antagonist initiated at day 14 significantly increased the number of microglia-engulfed neurons compared with vehicle treatment. Data are presented as mean ± s.e.m. n = 9 fields of view (FOVs) from 3 animals per group. Detailed statistics are provided in Supplementary Table1.

To test the contribution of noradrenergic neuronal activity in awake mice, we chemogenetically inhibited locus coeruleus (LC) noradrenergic neurons. LC inhibition during the active phase increased microglial phagocytosis **(Figure 4E-G)**. We next examined the β2-adrenergic receptor ADRB2 as a candidate mediator. Pharmacological activation of ADRB2 during the resting phase suppressed neuronal clearance, whereas inhibition during this phase enhanced phagocytic activity **(Figure 4H and 4I)**. Repeated daily ADRB2 blockade during the resting phase maintained enhanced clearance over the treatment period **(Figure 4J)**. Together, these interventions support a braking action of noradrenergic signaling on microglial engulfment and implicate ADRB2 in the regulation of neuronal clearance across the rest–activity cycle.

### Microglial ADRB2 deletion impairs clearance and blunts its light–dark regulation

Our pharmacological experiments identify ADRB2 signaling as a brake on microglial engulfment. Physiological changes in NE availability could therefore strengthen this restraint during the active phase and relieve it during rest. We next asked how sustained removal of this regulatory input affects clearance and its light–dark organization. We generated microglia-specific *Adrb2* conditional knockout mice (*Adrb2* cKO). RNAscope confirmed efficient deletion in microglia, while morphological analysis showed increased process complexity without altered microglial abundance **(Figure 5A; Figure S10A-G)**.

**Figure 5.**
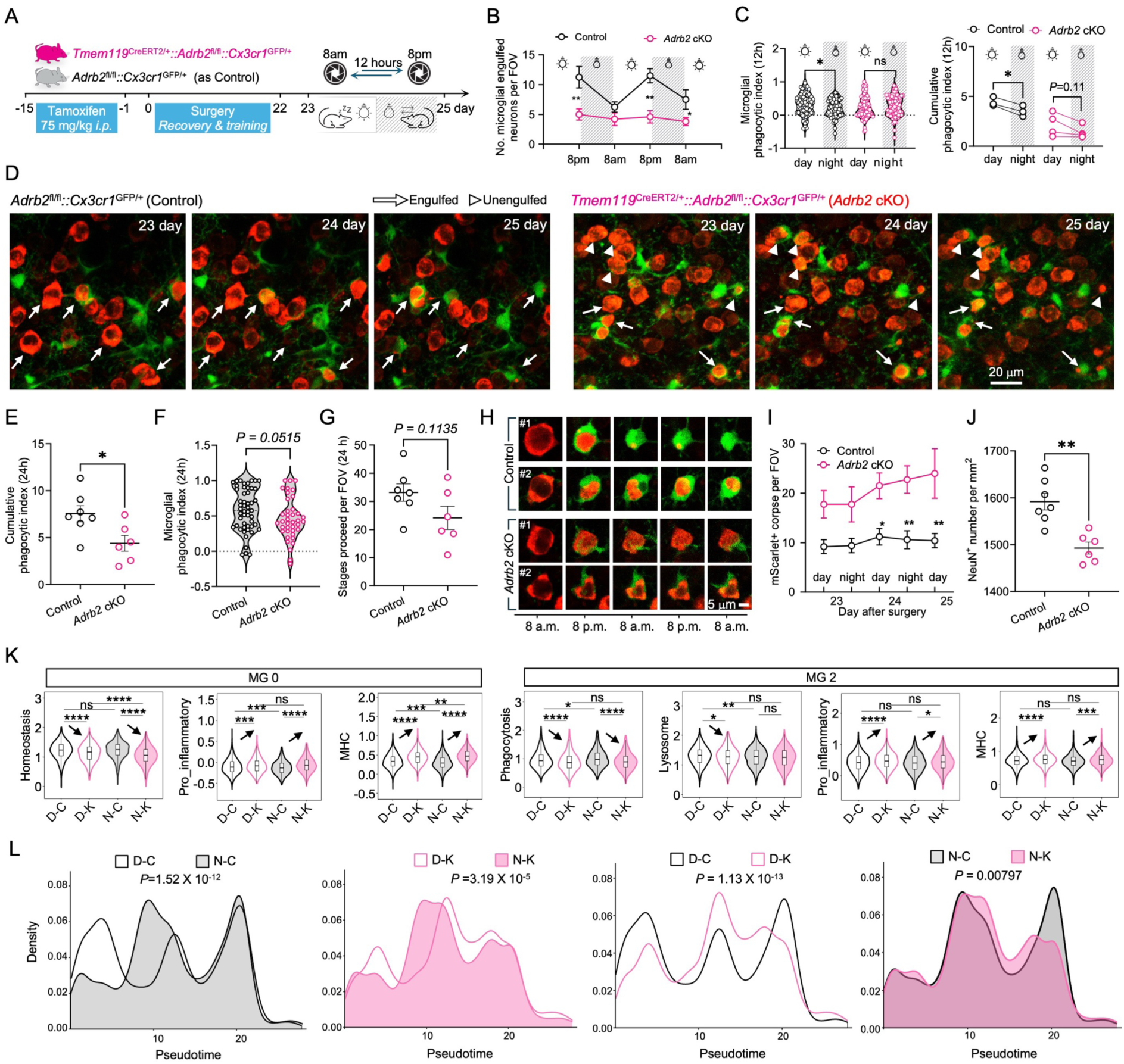
Conditional *Adrb2* deletion reduces phagocytic capacity and attenuate day-night modulation of microglial behavior. **(A)** Experimental design for in vivo two-photon imaging of microglial phagocytosis in conditional *Adrb2* knockout (*Adrb2* cKO) mice. **(B)** Quantification of engulfed neurons across daytime and nighttime imaging sessions. *Adrb2* cKO markedly reduced overall engulfment and attenuated day-night differences. Data represent mean ± s.e.m. n=4 animals for Cx3cr1^GFP/+^ group, n=5 animals for *Adrb2* cKO group. **(C)** Phagocytic capacity per microglia (left) and total engulfment events per field of view (FOV; right) in Cx3cr1^GFP/+^ and *Adrb2* cKO mice, showing no significant difference between day and night in *Adrb2* cKO mice. Data represent mean ± s.e.m. n=3 animals for Cx3cr1^GFP/+^ group, n=4 animals for *Adrb2* cKO group (c, right), n represents individual event pooled from (c) right. **(D)** Representative consecutive images of neuronal phagocytosis in Cx3cr1^GFP/+^ control and *Adrb2* cKO mice. White arrows indicate successfully phagocytosed neurons, whereas white arrowheads mark neuronal corpses that failed to be cleared. Scale bars, 20 μm. **(E-G)** Quantification of cumulative phagocytic index (E), microglial phagocytic index (F) and stages proceed (G), demonstrating reduced phagocytosis activity in *Adrb2* cKO mice compared with controls. Data represent mean ± s.e.m. For (E, G), n=7 animals for Cx3cr1^GFP/+^ group, n=6 animals for *Adrb2* cKO group. For (F), n represents individual event pooled from (E). **(H)** Representative high-resolution images showing interactions between individual microglia and degenerating neurons. Scale bars, 5 μm. **(I)** Quantification of uncleared neuronal corpses, revealing significant accumulation in *Adrb2* cKO mice compared with controls. Data represent mean ± s.e.m. n=5 animals for Cx3cr1^GFP/+^ group, n=4 animals for *Adrb2* cKO group. **(J)** Quantification of NeuN^+^ density shows *Adrb2*-cKO accelerate neuronal loss in TDP-43 model. Data represent mean ± s.e.m. n=7 animals for Cx3cr1^GFP/+^ group, n=6 animals for *Adrb2* cKO group. **(K)** Violin plots showing functional signature scores of MG 0 and MG 2 in Adrb2 cKO (pink) and control (grey) during the day and night. *, *P*<0.05, **, *P*<0.01, ***, *P*<0.001, ****, *P*<0.0001, one-way ANOVA followed by Tukey’s post-hoc test. **(L)** Kernel density plots showing the distribution of pseudotime across total microglia from different experimental groups. Pseudotime was inferred using Monocle3 trajectory analysis. Statistical significance was assessed using two-sided Wilcoxon rank-sum tests. D-C: day-control; D-K: day-*Adrb2* cKO; N-C: night-control; N-K: night-*Adrb2* cKO. Detailed statistics are provided in Supplementary Table1.

Despite the enhancement of engulfment following pharmacological inhibition, longitudinal imaging in the cytoTDP-43 model showed that sustained ADRB2 deletion reduced neuronal engulfment during both light and dark phases **(Figure 5B)**. Deletion also blunted the difference between phases and reduced both single-cell and cumulative clearance **(Figure 5C)**. ADRB2-deficient microglia continued to contact degenerating neurons but progressed inefficiently through RAID, resulting in prolonged interactions and accumulation of uncleared remnants **(Figure 5D-I)**. This impairment was accompanied by greater neuronal loss in the cKO cortex **(Figure 5J; Figure S10H)**.

To characterize molecular changes accompanying impaired clearance, we compared microglial transcriptional profiles from control and cKO mice sampled during the day and night. The four groups—day-control (D-C), day-cKO (D-K), night-control (N-C) and night-cKO (N-K)— allowed genotype-associated changes to be examined within each phase. Using the microglial populations defined above, we found that *Adrb2* deletion reduced the relative representation of homeostatic MG0 and DAM-like MG2 while expanding inflammatory-primed MG1 and interferon-responsive MG5 during both phases **(Figure S11A-C)**.

Within each phase, cKO microglia showed greater enrichment of inflammatory, interferon-responsive, chemotactic and antigen-presentation programs and reduced homeostatic and phagocytosis-associated signatures relative to controls **(Figure S11D and 11E)**. MG0 and MG2 showed particularly pronounced responses to deletion. Gene Ontology analysis identified alterations in phagocytic and metabolic programs alongside enrichment of inflammatory and interferon-related pathways **(Figure S11F and S11G)**. Signature analysis further identified changes in phagocytic, lysosomal, interferon and antigen-presentation programs within MG0 and MG2 **(Figure 5K; Figure S12)**. Pseudotime analysis was consistent with an altered organization of microglial transcriptional states under ADRB2 deficiency, with a bias towards inflammatory rather than phagocytosis-associated programs **(Figure 5L)**.

Given the metabolic demands of corpse clearance^15,31^, we also examined metabolic gene signatures. Control microglia showed phase-associated differences, with greater resting-phase enrichment of programs related to amino-acid metabolism, lipid remodeling, carbohydrate utilization, energy metabolism and lysosomal function. ADRB2-deficient microglia exhibited reduced enrichment of these programs, particularly within MG2 **(Figure S13)**. These findings associate the clearance impairment with changes in microglial population composition and phagocytic, inflammatory and metabolic gene programs.

Together, these results distinguish the acute braking action of ADRB2 signaling from the consequences of sustained receptor loss. Although pharmacological inhibition enhances engulfment, microglial ADRB2 deletion compromises clearance and blunts its light–dark regulation, accompanied by altered transcriptional profiles. The findings support a role for ADRB2 both in noradrenergic restraint of engulfment and in the maintenance of effective, phase-dependent clearance.

## Discussion

Our findings identify microglial removal of degenerating neurons as a cellular maintenance process coordinated with the host’s rest–activity cycle. Longitudinal imaging resolved an ordered sequence of microglial engagement and corpse removal, while behavior-guided transcriptomics associated engulfment with distinct molecular programs. Clearance was enhanced during the resting phase, followed the shifted light–dark schedule after cycle inversion and showed reduced phase dependence following sleep disruption. Noradrenergic interventions further identified NE–ADRB2 signaling as a brake on engulfment. Sustained microglial ADRB2 deletion nevertheless impaired clearance and blunted its light–dark variation, indicating that the consequences of receptor loss extend beyond relief of acute inhibition.

### Microglial phagocytosis is essential for efficient neuronal corpse clearance

PLX5622-mediated depletion distinguished neuronal degeneration from subsequent corpse removal. Neuronal shrinkage and structural collapse continued in the absence of microglia, whereas cellular remnants persisted until microglial repopulation. These observations establish a requirement for microglia in efficient corpse removal in this model. Depletion was also associated with greater neuronal loss, although this finding does not establish that defective clearance alone accounts for the worsening pathology. The results are consistent with evidence that microglia can support tissue recovery in TDP-43 proteinopathy^32^ and emphasize the importance of distinguishing their clearance functions from their broader effects on neuronal survival^33^.

### Linking microglial behavior to transcriptional state

Single-cell transcriptomic studies have revealed extensive microglial heterogeneity, but transcriptional state alone does not necessarily indicate what an individual microglial cell is doing in situ^34,35^. By selecting microglia according to observed engagement status, CBG-seq linked resting, activated but non-engulfing and engulfing cells to distinct molecular profiles. Activated but non-engulfing microglia remained transcriptionally distinguishable from cells observed engulfing neuronal material, demonstrating that activation and engulfment are not interchangeable descriptions of microglial behavior. Engulfment was associated with greater expression of lysosomal, metabolic and cargo-processing gene programs.

Mapping these behavior-defined signatures onto conventional single-cell datasets suggests that some heterogeneity within disease-associated microglial populations may reflect differences in target engagement and clearance. This provides a functional perspective on transcriptional diversity beyond a homeostatic-to-inflammatory axis. However, the sampled categories do not establish continuous transitions within individual cells, and the associated expression changes may precede engulfment, result from cargo uptake, or reflect both. The profiles therefore identify candidate programs associated with clearance rather than establish the molecular sequence controlling RAID progression.

### Microglial clearance is coordinated with host rest-activity state

Microglial inflammatory, phagocytic and metabolic functions have been reported to vary across the day^36–41^. Our findings extend this temporal organization to the removal of degenerating neuronal somata. Clearance was enhanced during the resting phase, its pattern reversed relative to clock time after light–dark cycle inversion, and the difference between phases was blunted by sleep disruption. Together, these observations link clearance to the organization of rest and activity, while leaving open the respective contributions of sleep, arousal and circadian regulation.

Why might clearance be enhanced during rest? Engulfment and degradation of a neuronal soma involve cytoskeletal remodeling, membrane trafficking and lysosomal processing^42,43^. In our dataset, resting-phase microglia showed greater enrichment of metabolic, lysosomal and phagocytosis-associated gene programs, whereas active-phase profiles showed stronger inflammatory and stress-responsive signatures. These associations suggest that rest may provide conditions favorable for cellular clearance, although gene-expression signatures alone do not establish increased metabolic flux or lysosomal activity.

The physiological advantage of this timing remains unresolved. One possibility is that rest provides conditions favorable for the sustained cellular activity required for engulfment and degradation. Whether this temporal organization also limits disruption of neighboring circuits will require simultaneous measurements of neuronal activity and microglial clearance. Our experiments do not establish a direct trade-off between neuronal information processing and corpse removal.

### Noradrenergic restraint and sustained ADRB2-dependent regulation of clearance

Our intervention experiments support an acute braking action of NE–ADRB2 signaling on microglial engulfment. This interpretation is consistent with evidence that NE suppresses microglial process dynamics, and that β-adrenergic stimulation can inhibit phagocytosis^18,19,44^. Reduced noradrenergic input during rest could therefore relieve this restraint and contribute to enhanced clearance. Microglial ADRB2 provides a potential route through which changes in NE availability can link cellular behavior to the host’s physiological state.

Sustained microglial ADRB2 deletion, however, impaired RAID progression, reduced clearance and favored inflammatory and interferon-associated transcriptional profiles. These outcomes differ from the enhancement observed following pharmacological inhibition and indicate that receptor deletion cannot be treated as equivalent to prolonged relief of acute restraint. Differences in duration, completeness and cellular specificity of the perturbations, together with adaptation to receptor loss, could contribute to the divergent phenotypes.

The findings raise the possibility that responsiveness to changing NE availability helps maintain the coordination of clearance with host rest and activity. However, deletion removes both tonic and time-varying ADRB2 input; it does not establish that loss of physiological signaling fluctuations causes clearance failure. β2-adrenergic signaling can regulate inflammatory and metabolic programs in myeloid cells^45–47^, providing candidate mechanisms through which sustained receptor loss might affect microglial function. The accompanying transcriptional changes are consistent with altered cellular state, but direct functional measurements and targeted rescue experiments will be needed to determine whether these changes mediate the clearance deficit.

### Rest-associated clearance as a component of brain maintenance

Sleep-associated changes in fluid dynamics have been implicated in the redistribution and removal of soluble metabolites and pathogenic proteins^48,49^. Our findings extend the question of rest-associated maintenance to the removal of entire degenerating neurons. Fluid-mediated transport and microglial corpse clearance operate at different biological scales: the former redistributes soluble material, whereas the latter involves cellular recognition, encapsulation and degradation. Their association with resting states raises the possibility that complementary maintenance processes are organized around host physiology, although their coordination was not directly examined here.

This perspective may be relevant to neurodegenerative disease. Sleep disruption commonly accompanies neurodegeneration and has been linked to changes in pathological protein clearance and microglial function^50–56^. Pathological changes in the locus coeruleus– noradrenergic system in several neurodegenerative disorders^57–59^, could also alter arousal and the signals received by microglia. Notably, pharmacologically increasing noradrenergic transmission has produced limited cognitive or biomarker effects and has not been shown to slow neurodegenerative disease progression^60,61^. These observations motivate consideration of both the level and temporal pattern of noradrenergic signaling when investigating its contribution to disease. Whether preserving or restoring state-dependent signaling improves neuronal clearance or disease outcomes remains unknown.

### Limitations

Several limitations should be considered. First, the model is driven by aberrant cytosolic TDP-43 expression and does not reproduce the full complexity of human TDP-43 proteinopathies or other neurodegenerative diseases. Use of mouse TDP-43 avoids introducing a human protein sequence but does not remove the limitations of overexpression or the nuclear localization sequence mutation. Validation in additional disease models, aging and human tissues will be necessary to establish generality.

Second, light–dark inversion and sleep disruption do not fully separate the contributions of sleep itself, arousal-related behavior, environmental timing and endogenous circadian mechanisms. Stress or movement associated with sleep-disruption procedures may also contribute to the observed effects. Microglia possess intrinsic molecular clocks^62^; experiments under constant environmental conditions, direct sleep-state measurements and manipulation of core clock genes could help determine how these mechanisms interact with NE–ADRB2 signaling.

Third, CBG-seq associates engagement categories with transcriptional profiles but does not establish which molecular programs causally control RAID progression. Longitudinal molecular measurements and targeted manipulation of candidate pathways will be needed to distinguish changes that enable engulfment from those induced by cargo uptake. Direct measurements of metabolic and lysosomal function are also required to test the functional implications of the transcriptional signatures.

Fourth, the mechanism underlying impaired clearance after sustained ADRB2 deletion remains unresolved. The pharmacological and genetic interventions differ in more than duration, and their comparison does not isolate the role of physiological signaling fluctuations. Moreover, reduced overall clearance could itself diminish the observed light–dark contrast. Distinguishing altered temporal regulation from a general loss of clearance capacity will require analysis of phase-dependent effects alongside overall activity and experiments that manipulate the timing of signaling while controlling its overall level.

Finally, the relationship between clearance and neuronal survival remains incompletely resolved. Greater neuronal loss following depletion or ADRB2 deletion may involve microglial functions beyond corpse removal. Experiments that selectively alter clearance, together with measurements of neighboring neuronal activity and survival, will be needed to establish its contribution to tissue protection.

Together, our findings show that microglial removal of degenerating neurons is enhanced during the host resting phase and regulated by noradrenergic signaling. NE–ADRB2 signaling restrains acute engulfment, whereas sustained microglial receptor loss impairs clearance and blunts its light–dark variation. These results link cellular corpse removal to host physiological state and provide a framework for investigating how the timing of microglial clearance contributes to brain maintenance.

## Acknowledgments

This work was supported by the National Key R&D Program of China (Grant No. 2023YFA1800100 to Y.U.L.), the National Natural Science Foundation of China (Grant No. 82071188 to Y.U.L.), the Science and Technology Projects in Guangzhou (Grant No. 2024A03J1256 to Y.U.L.), and Science and Technology Projects in Guangzhou (Grant No. 2025A03J4156 and 2024A04J3932 to Z.X.).

## Author contributions

Z.X., B.M. and Y.U.L. prepared the manuscript with help from other authors. Z.X., B.M. and Y.U.L. conceived and designed this research. Z.X. and B.M. performed the experiments with help from M.L., J.C., and L.H.. Z.X., B.M. and M.L. performed data analysis. Z.X. and Y.U.L. initiated and supervised the project. Z.X., B.M., M.L., and Y.U.L. are the guarantors of this work and, as such, had full access to all the data in the study and take responsibility for the integrity of the data and the accuracy of the data analysis.

## Data availability

The source data underlying all Figures are provided with this paper. Any additional information required to reproduce the results reported in this study is available from the corresponding author upon reasonable request.

## Code availability

All code used in the analysis of scRNA-seq datasets and generation of Figures are available upon reasonable request.

---

**Figure S1.**
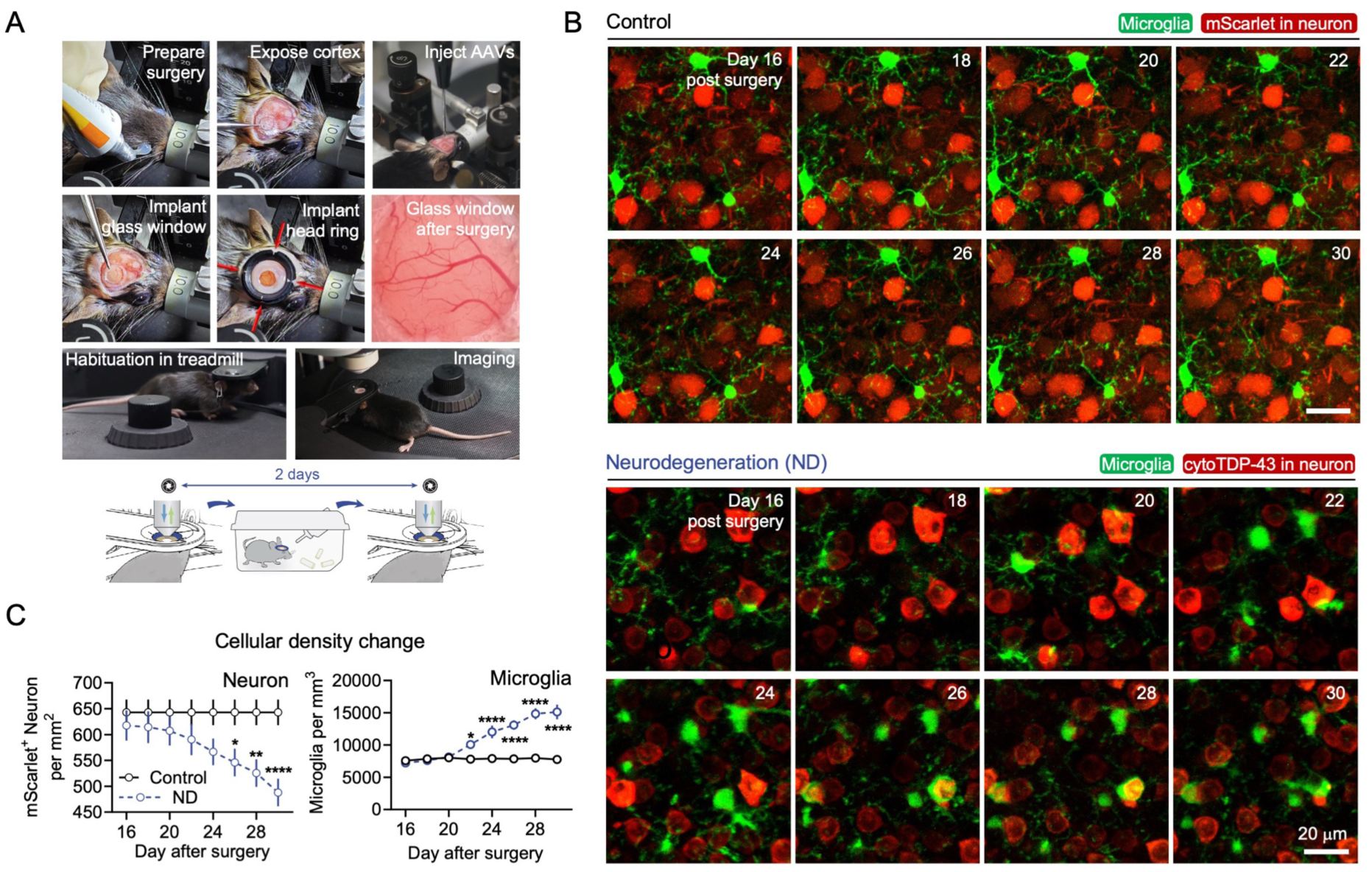
Chronic awake two-photon imaging platform for longitudinal tracking of microglial–neuronal interactions during neurodegeneration. **(A)** Workflow of the experimental pipeline integrating AAV-injection, chronic cranial window implantation, and awake two-photon imaging. Representative images show sequential steps of surgery, including cortical exposure, viral injection, glass window implantation, and head-ring fixation. Following recovery and treadmill habituation, mice were repeatedly imaged under awake conditions, enabling longitudinal tracking of microglial and neuronal dynamics within the same cortical regions over time. **(B)** Repeated two-photon imaging at 2-day intervals of AAV-labeled neurons (mScarlet) and GFP-labeled microglia under control or neurodegenerative conditions. In controls, neuronal morphology and microglial architecture remained stable. In TDP-43 group, neurons exhibited a stereotyped degenerative trajectory characterized by progressive fluorescence accumulation, somatic shrinkage and eventual signal extinction, followed by rapid microglial envelopment and phagocytic clearance. Scale bar, 20 μm. **(C)** Quantitative analysis of neuronal and microglial density dynamics across imaging sessions. n=5 animals for each group. Detailed statistics are provided in Supplementary Table1.

**Figure S2.**
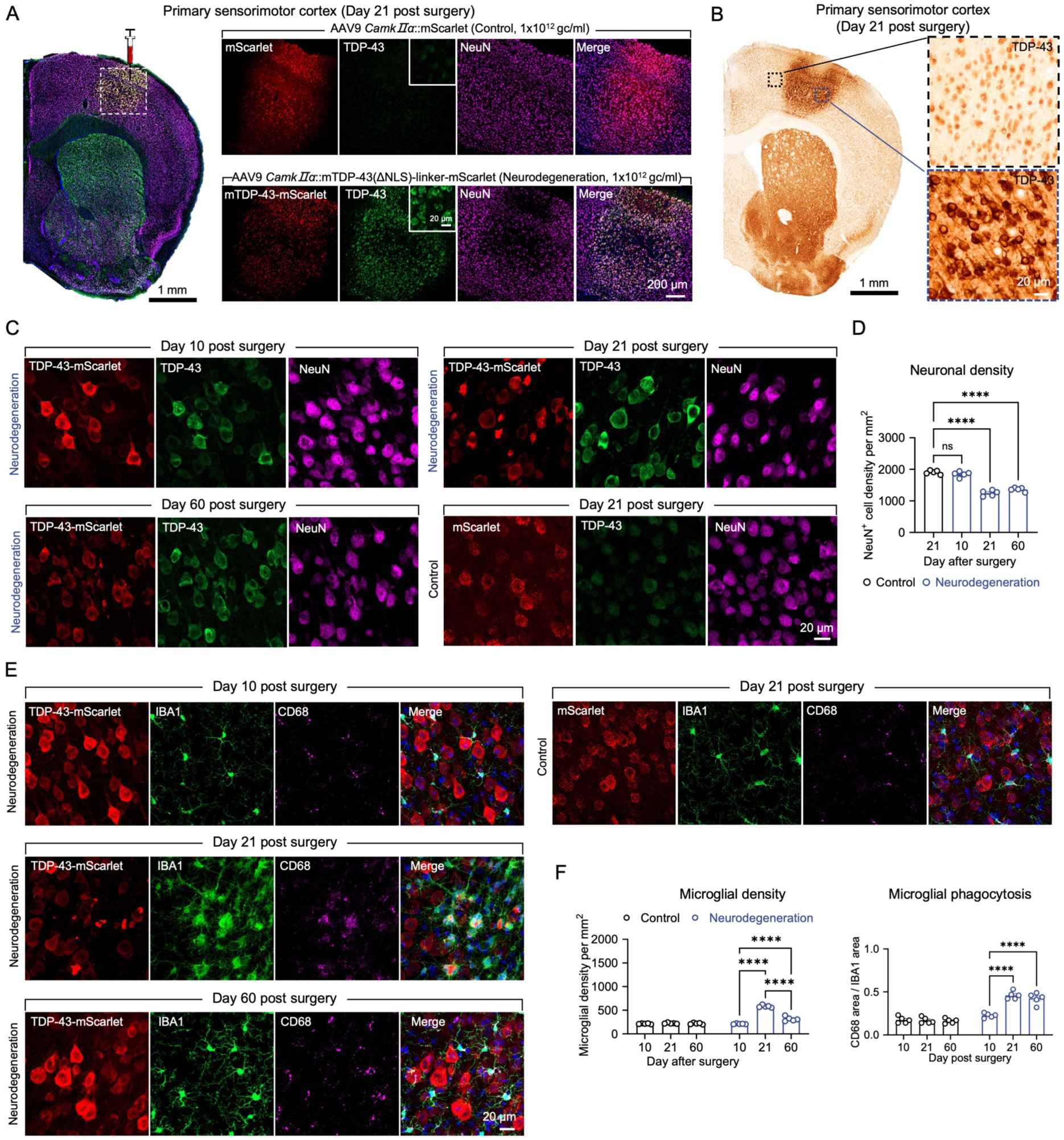
Overexpression of cytosolic TDP-43 in mouse primary motor cortex induces a neurodegeneration model. **(A, B)** Brain sections showing TDP-43 expression in the primary motor cortex (M1) 21 days after AAV injection, visualized by immunofluorescence (A) and immunohistochemistry (B). **(C)** Time-course analysis showing that significant neuronal loss emerges by 21 days post-injection. Scale bars, 20 μm. **(D)** Quantification of NeuN⁺ neuronal density at different time points post-injection. Data represent mean ± s.e.m. n = 5 animals for each group. **(E)** Representative immunofluorescence images showing microglial density and CD68 expression in control and TDP-43–overexpressing groups at three time points. Scale bars, 20 μm. **(F)** Quantification of microglial density and CD68 levels across time, demonstrating progressive microglial proliferation and enhanced phagocytic activation in the TDP-43 group. Data represent mean ± s.e.m. n=5 animals for each group. Detailed statistics are provided in Supplementary Table1.

**Figure S3.**
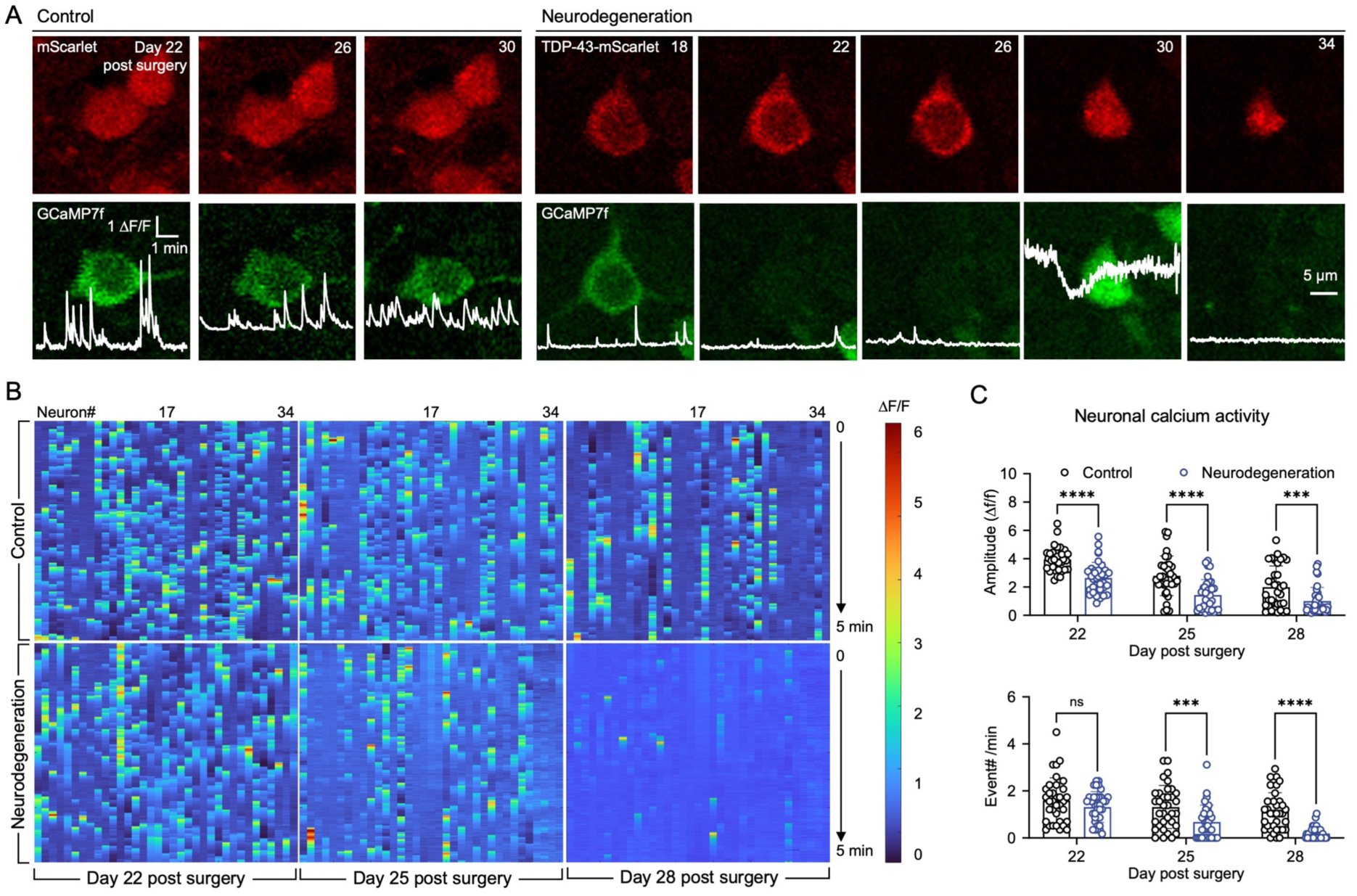
Two-photon calcium imaging reveals impaired neuronal activity in the cytoTDP43 model. **(A)** Representative images of mScarlet-labeled neurons expressing GCaMP7f and corresponding calcium traces in control (left) and cytoTDP43 (right) groups. Neurons in the control group exhibit stable calcium events frequency, whereas TDP43 neurons show markedly reduced activity. Scale bars, 5 μm. **(B)** Heatmap illustrating progressive decline in neuronal calcium activity from day 22 to day 28 in the cytoTDP43 group. **(C)** Quantification of calcium event amplitude and frequency across three time points in control versus cytoTDP43 groups. Data represent mean ± s.e.m. n=35 neurons from 3 animals for each group. Detailed statistics are provided in Supplementary Table1.

**Figure S4.**
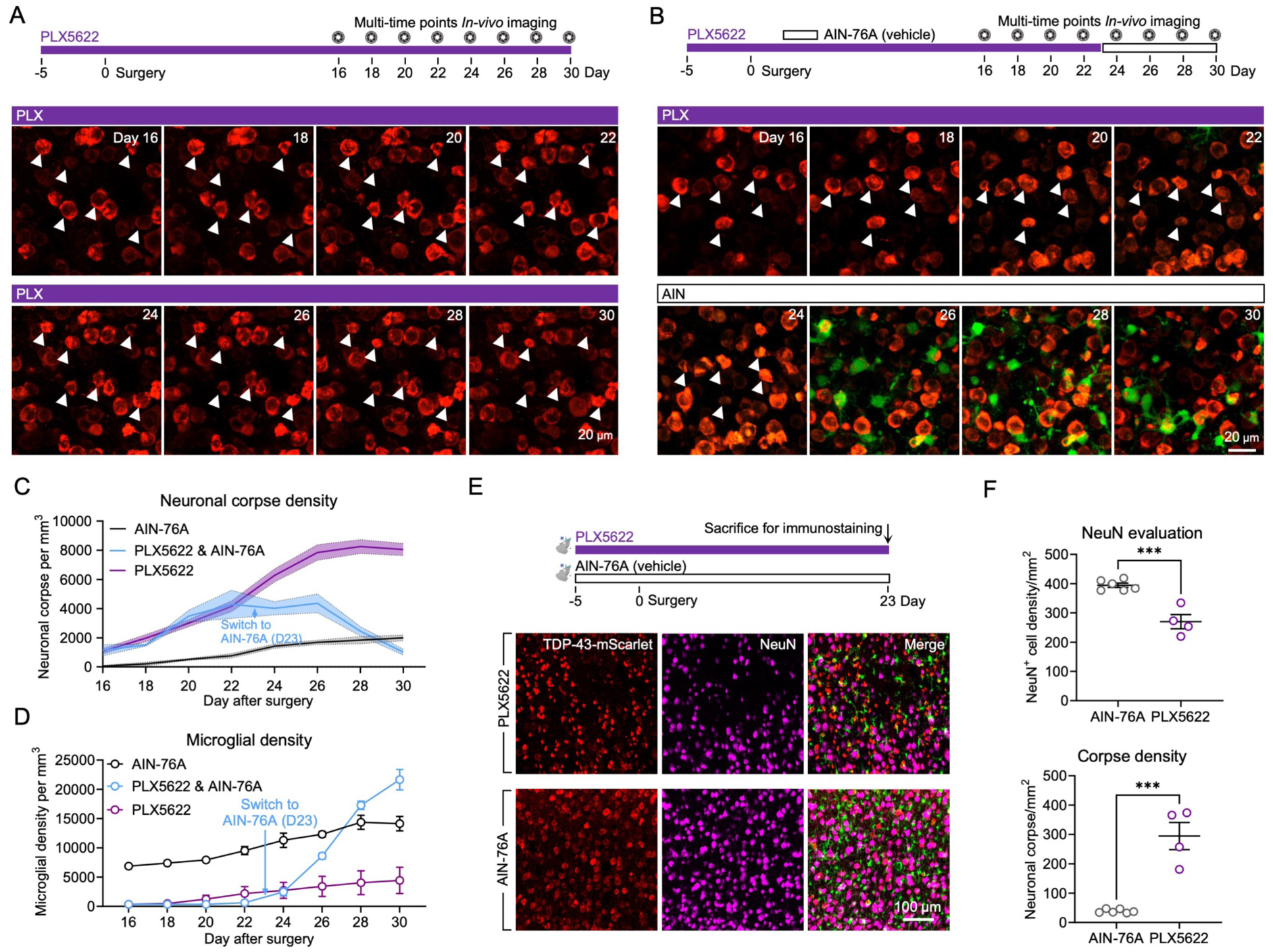
Sustained PLX5622-mediated microglial depletion exacerbates cytoTDP43 pathology. **(A, B)** Experimental timeline showing continuous PLX5622 treatment starting 5 days prior to TDP43-AAV injection, with longitudinal two-photon imaging initiated 16 days after surgery. Representative two-photon images show persistent neuronal corpses (white arrowheads) following microglial depletion and their subsequent clearance during microglial repopulation after PLX5622 withdrawal. Scale bars, 20 μm. **(C)** Quantification demonstrating increased neuronal corpse accumulation during microglial depletion and reduction following microglial repopulation. Trend lines represent the mean ± s.e.m. n=4 animals for AIN-76A group; n=3 animals for PLX5622 and PLX5622&AIN-76A group. **(D)** Microglial density measurements under different conditions. Sustained PLX5622 treatment maintained low microglial density, while withdrawal at day 23 triggered robust microglial proliferation. Trend lines represent the mean ± s.e.m. n=4 animals for AIN-76A group; n=3 animals for PLX5622 and PLX5622&AIN-76A group. **(E)** Experimental design for pathological analyses following microglial depletion prior to TDP43-AAV injection. Confocal images showing elevated neuronal corpse density (mScarlet⁺/NeuN⁻ cells) and reduced NeuN⁺ neuronal density in the PLX5622-treated group. Scale bars, 100 μm. **(F)** Quantification of NeuN⁺ neuronal density and neuronal corpse density in AIN-76A versus PLX5622-treated groups. Data represent mean ± s.e.m. n=6 animals for AIN-76A group, n=4 animals for PLX5622 group. Detailed statistics are provided in Supplementary Table1.

**Figure S5.**
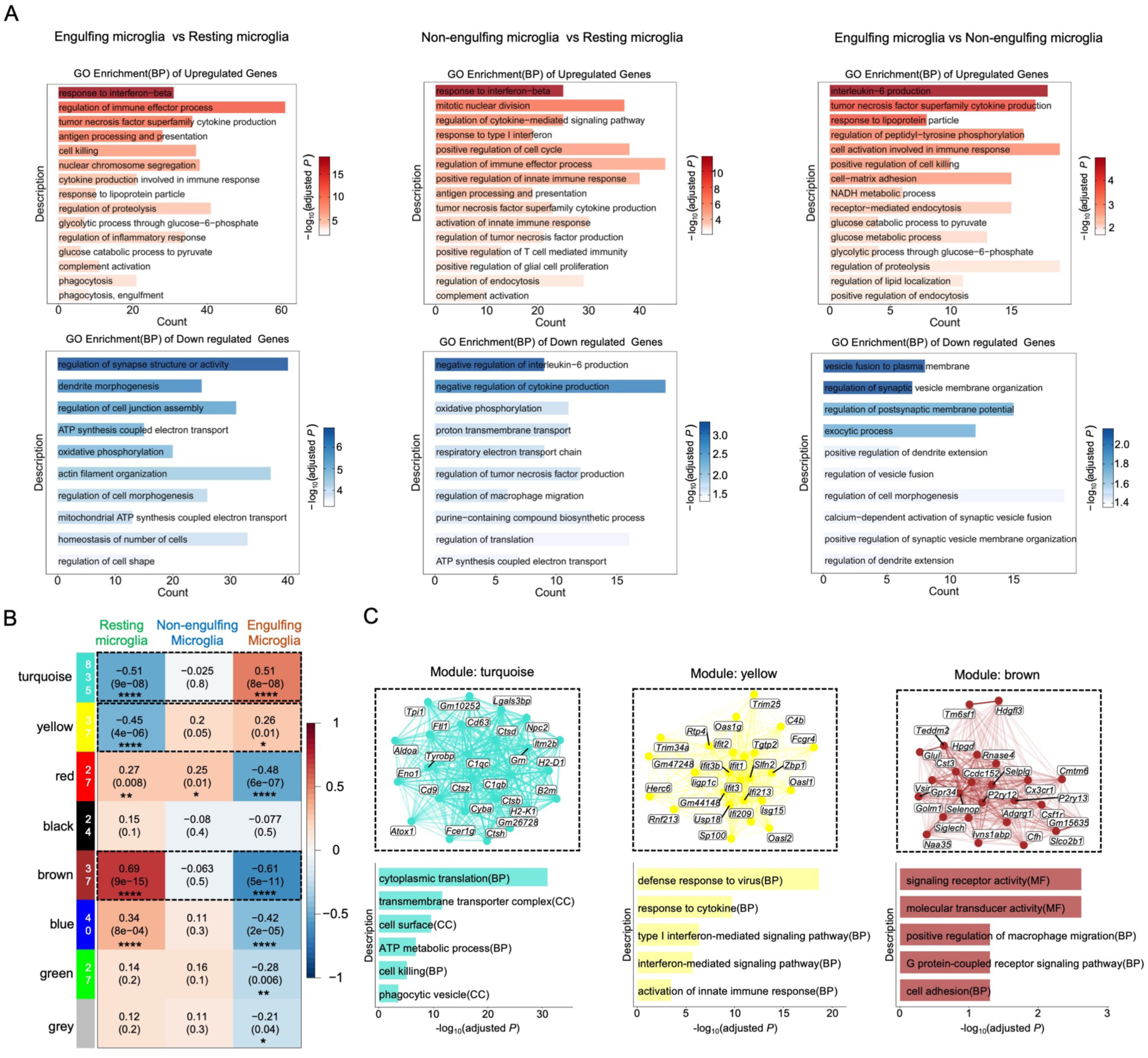
Transcriptional states associated with imaging-guided microglial states. **(A)** GO biological process enrichment analysis of the significantly upregulated genes (top) and downregulated genes (bottom). **(B)** WGCNA identified eight gene co-expression modules, and their associations with resting, non-engulfing and engulfing microglia were assessed by Pearson correlation. Colors indicate Pearson correlation coefficients, with corresponding *P* values shown in parentheses. **(C)** The top 25 hub genes ranked by module eigengene-based connectivity (kME). Functional annotations were based on GO enrichment analysis. BP, biological process; MF, molecular function; CC, cellular component. *, *P* <0.05, **, *P*<0.01, ***, *P* <0.001, ****, *P*<0.0001.

**Figure S6.**
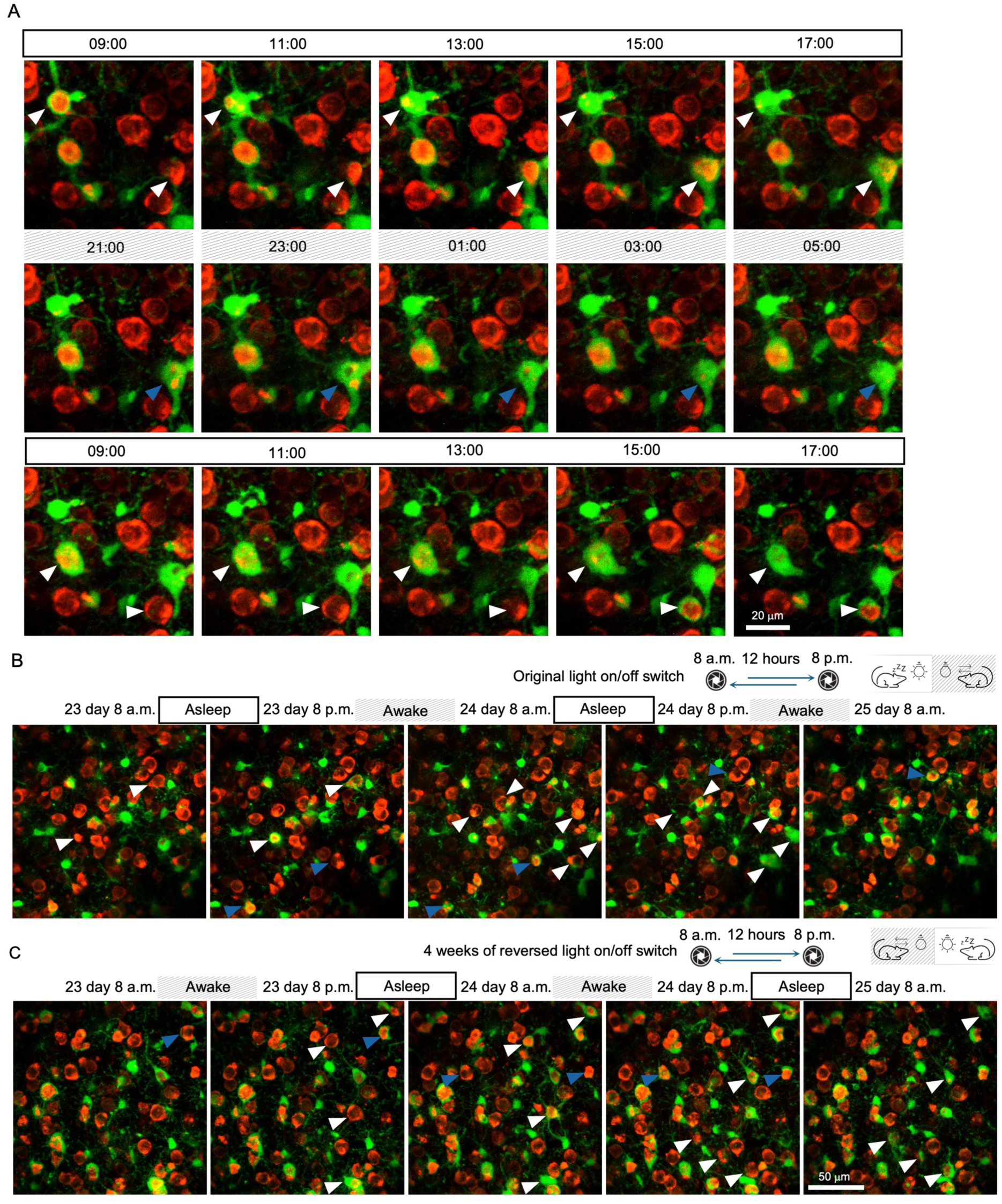
In vivo live imaging of microglial phagocytic clearance of degenerated neurons across day and night cycle. **(A)** Two hours interval image sessions show microglial phagocytosis enhanced in day (white arrowheads), decreased in night (blue arrowheads). Scale bar, 20 μm. **(B)** Representative imaging shows microglial phagocytosis enhanced in day in normal circadian (white arrowheads), decreased in night (blue arrowheads). **(C)** Day-night differences were inverted under the reversed circadian cycle. Scale bar, 50 μm.

**Figure S7.**
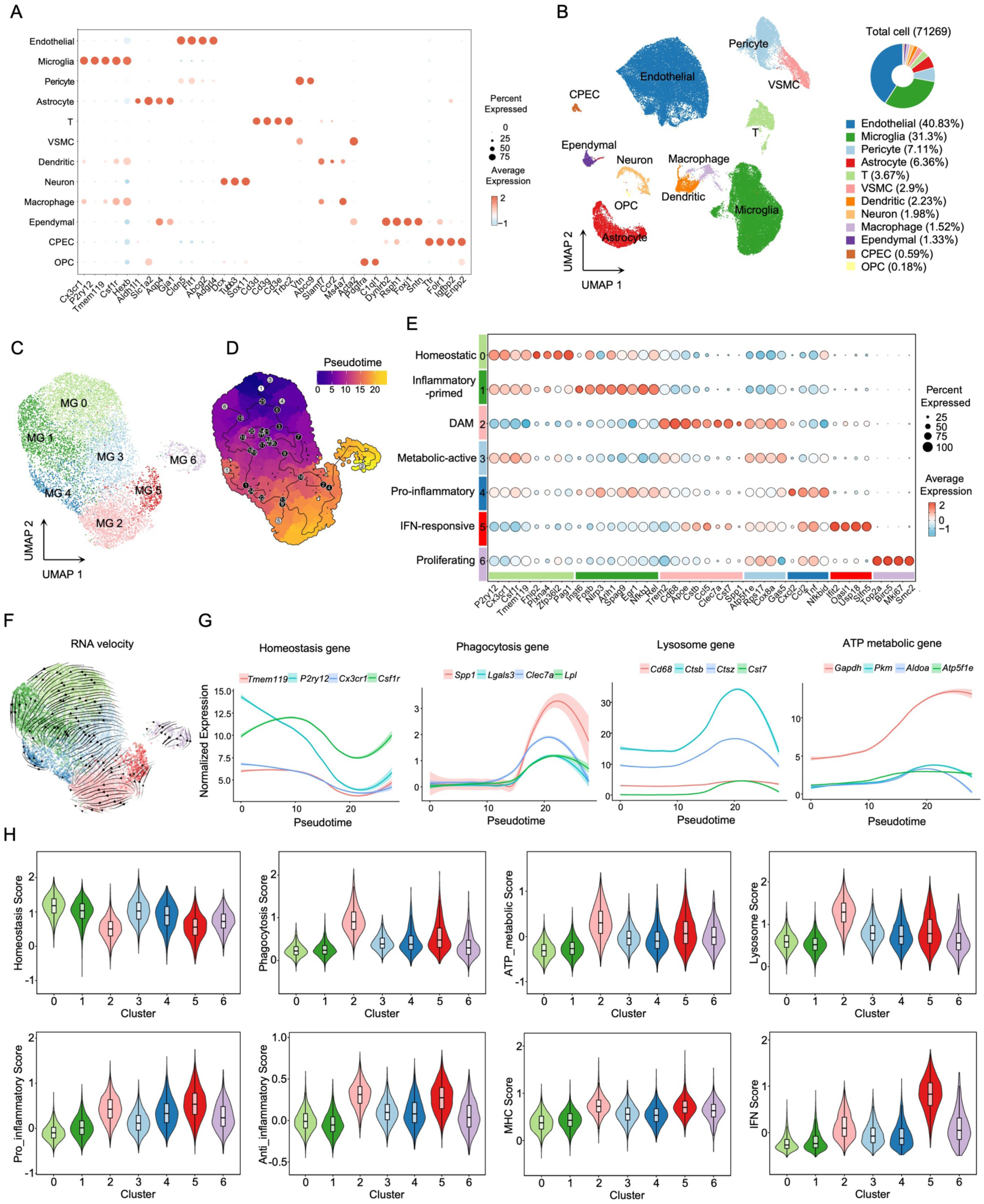
Single-cell RNA-seq analysis of cytoTDP-43 induced neurodegenerative cortex across day and night. **(A)** Canonical marker-gene expression across major cell types. **(B)** UMAP visualization of all 71,269 cells colored by cell type (left) and proportional cell-type composition (right). **(C)** UMAP visualization of microglial subclusters across the two phases. **(D)** Monocle3 pseudotime trajectory of microglia. **(E)** Marker-gene expression defining MG0, homeostatic; MG1, inflammatory-primed; MG2, DAM-like/engulfment-associated; MG3, metabolically active; MG4, pro-inflammatory; MG5, interferon-responsive; and MG6, proliferative populations. **(F)** scVelo RNA-velocity streamlines projected onto the UMAP embedding. **(G)** Representative homeostatic, phagocytic, lysosomal and ATP-metabolic genes plotted along Monocle3-inferred pseudotime. Lines indicate smoothed expression and shaded regions indicate 95% confidence intervals. **(H)** Functional signature scores across MG0–MG6 calculated using AddModuleScore.

**Figure S8.**
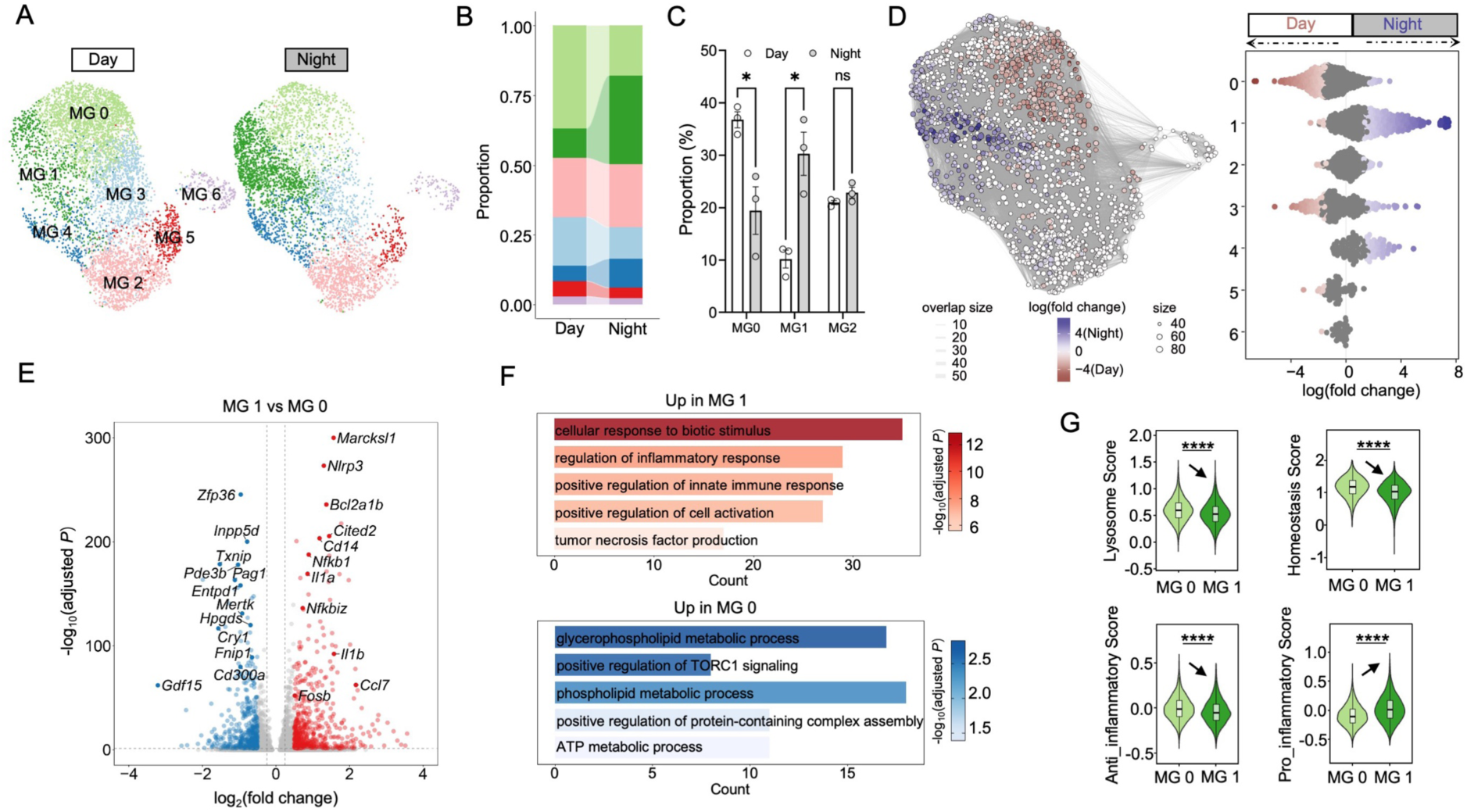
Day-night remodeling of microglial states during cytoTDP-43 induced neurodegeneration. **(A)** UMAP visualization of scRNA-seq profiles from neurodegenerative microglia collected during the day and night, colored by microglial subclusters. **(B)** Bar plot showing the proportion of microglial subclusters across day and night conditions. **(C)** Bar plot showing the proportion of MG0-MG2 in day and night. n=3 mice for each group. **(D)** Graphical representation of the neighborhood analysis using Milo (left). Relative enrichment of the microglia neighborhood clusters in day and night (right). **(E)** Volcano plots showing DEGs between MG1 versus MG0. **(F)** GO biological process enrichment analysis of significantly upregulated genes(top) and downregulated (bottom) in MG1 versus MG0. **(G)** Violin plots showing functional signature scores for phagocytosis, lysosome, homeostasis, pro-inflammatory, and anti-inflammatory programs in MG1 and MG0 microglia. Scores were calculated using AddModuleScore. ****, p < 0.0001; unpaired *t*-test (two-tailed). Detailed statistics are provided in Supplementary Table1.

**Figure S9.**
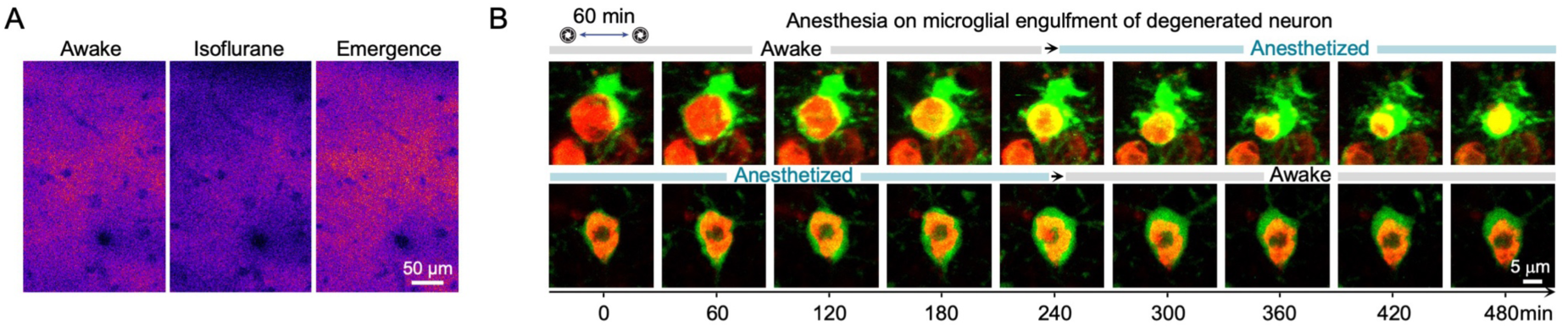
Anesthesia enhances microglial engulfment activity during in vivo imaging. **(A)** Representative images showing reduced NE sensor signals under anesthesia. Scale bars, 50 μm. **(B)** Representative time-lapse images illustrating microglial behavior during transitions between awake and anesthetized states. Scale bars, 5 μm

**Figure S10.**
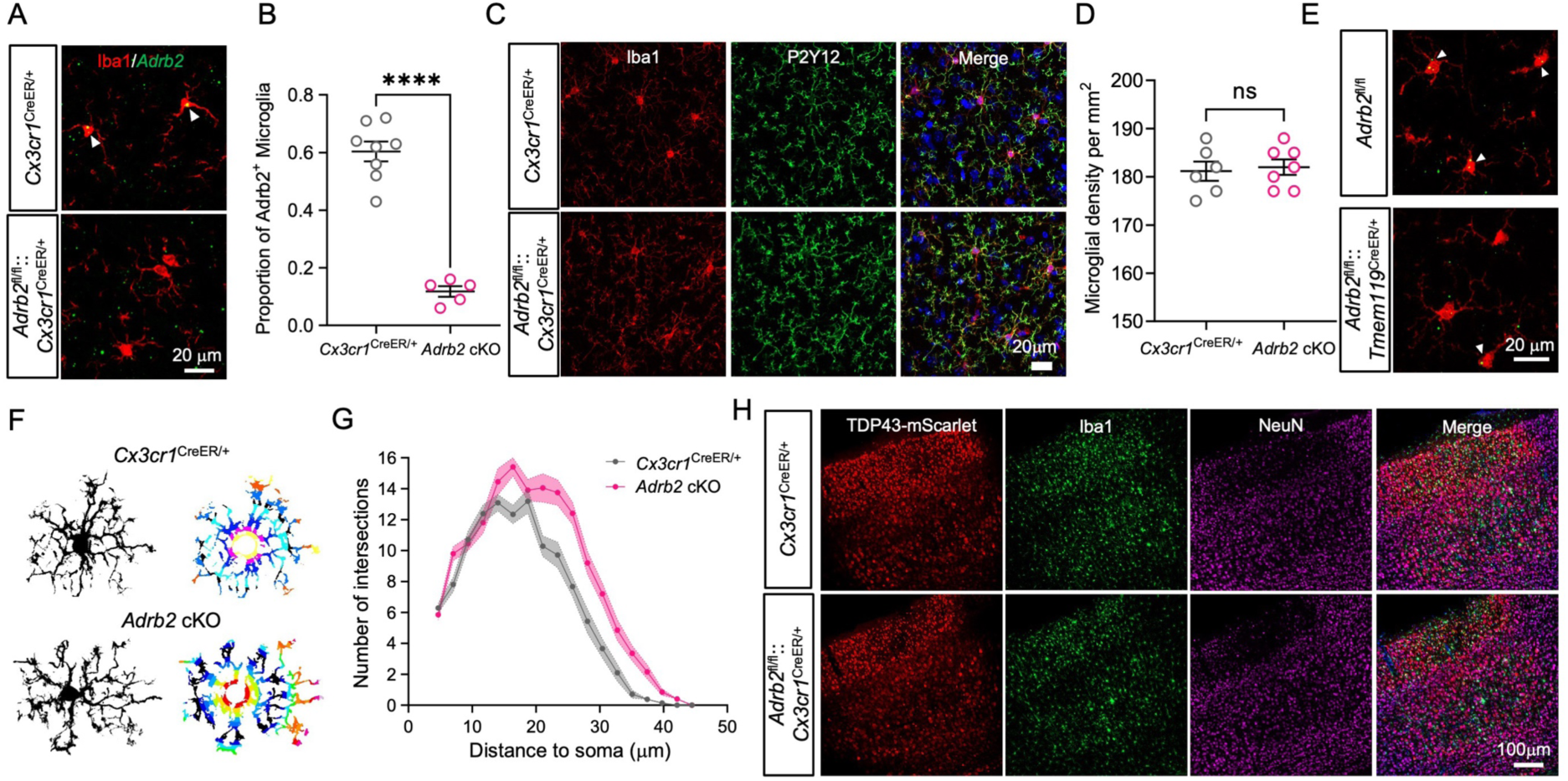
Characterization of microglial ADRB2 conditional knockout. **(A)** Fluorescence in situ hybridization (FISH) confirming efficient conditional knockout (cKO) of ADRB2 in microglia (arrowheads) driven by the Cx3cr1 promoter. Scale bar, 20 μm. **(B)** Quantification of ADRB2⁺ microglia showing a marked reduction in *Adrb2* cKO mice compared with controls. Data represent mean ± s.e.m. n=8 animals for Cx3cr1^CreER/+^ group, n=5 animals for *Adrb2* cKO group.**(C)** Representative images of microglial morphology labeled with Iba1 and P2ry12. Scale bar, 20 μm. **(D)** Quantification of microglial density in control and *Adrb2* cKO cortices, revealing no significant difference between groups. Data represent mean ± s.e.m. n=6 animals for Cx3cr1^CreER/+^ group, n=7 animals for *Adrb2* cKO group. **(E)** Fluorescence in situ hybridization (FISH) confirming efficient microglia-specific *Adrb2* deletion driven by the TMEM119 promoter (arrowheads). Scale bar, 20 μm. **(F-G)** Sholl analysis of microglial process complexity in control and *Adrb2* cKO mice, demonstrating increased process length and spatial coverage in ADRB2-deficient microglia. Trend lines represent the mean ± s.e.m. n=21 microglia from 5 animals for Cx3cr1^CreER/+^ group, n=20 microglia from 5 animals for *Adrb2* cKO group. **(H)** Representative images showing reduced NeuN⁺ neuronal density in Adrb2 cKO mice 21 days after TDP-43 injection. Scale bar, 20 μm. Detailed statistics are provided in Supplementary Table1.

**Figure S11.**
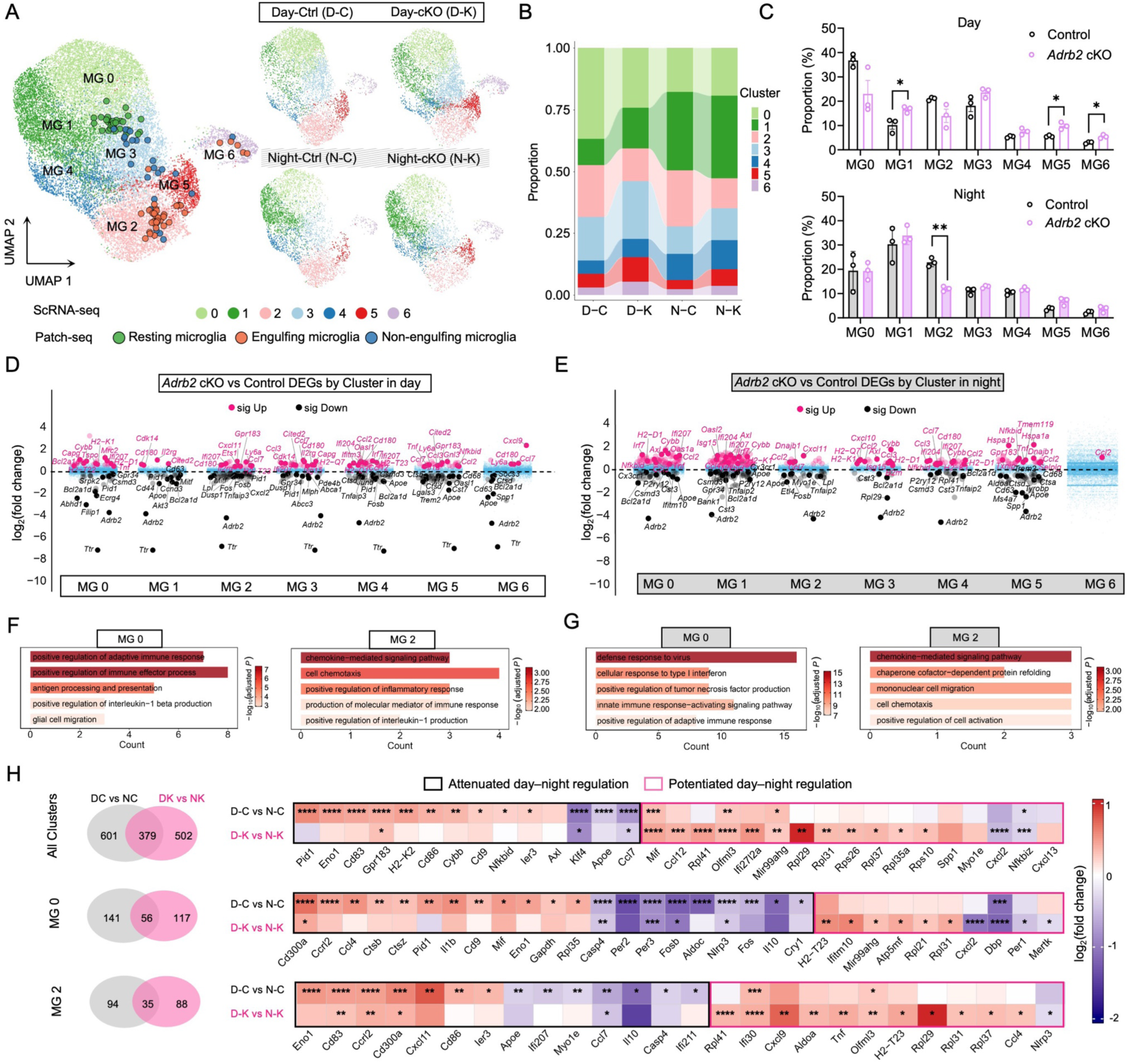
*Adrb2* deletion attenuates day–night transcriptional remodeling of microglia during cytoTDP-43-induced neurodegeneration. **(A)** UMAP visualization of microglial subclusters (MG0-MG6) identified by scRNA-seq (left), with transcriptional signatures of three microglial states (Resting, Engulfing, Non-engulfing) defined by SMART-seq mapped onto the scRNA-seq data. UMAPs (right) show microglial subcluster distribution in Day-Ctrl (D-C), Day-cKO (D-K), Night-Ctrl (N-C) and Night-cKO (N-K) groups. **(B)** Bar plot showing the proportion of each subclusters across groups. **(C)** Proportions of microglial subclusters (MG0–MG6) during the day and night in control and *Adrb2* cKO. n=3 mice for each group. **(D-E)** Cluster-wise volcano plots showing DEGs between *Adrb2* cKO versus control microglia within each microglia clusters (MG0-MG6) during day (D) and night (E). **(F-G)** GO biological process enrichment analysis of significantly upregulated genes in *Adrb2* cKO versus control in MG0 and MG2 during the day (F) and night (G). Detailed statistics are provided in Supplementary Table1. **(H)** Venn diagrams showing the overlap of day–night DEGs identified in control (D-C versus N-C) and *Adrb2* cKO (D-K versus N-K) microglia across total microglia, MG0 and MG2; Heatmaps display selected shared and condition-specific day–night DEGs. Differential expression significance was determined using Benjamini–Hochberg correction. *, adjusted *P*<0.05; **, adjusted *P*<0.01; ***, adjusted *P*<0.001; ****, adjusted *P*<0.0001.

**Figure S12.**
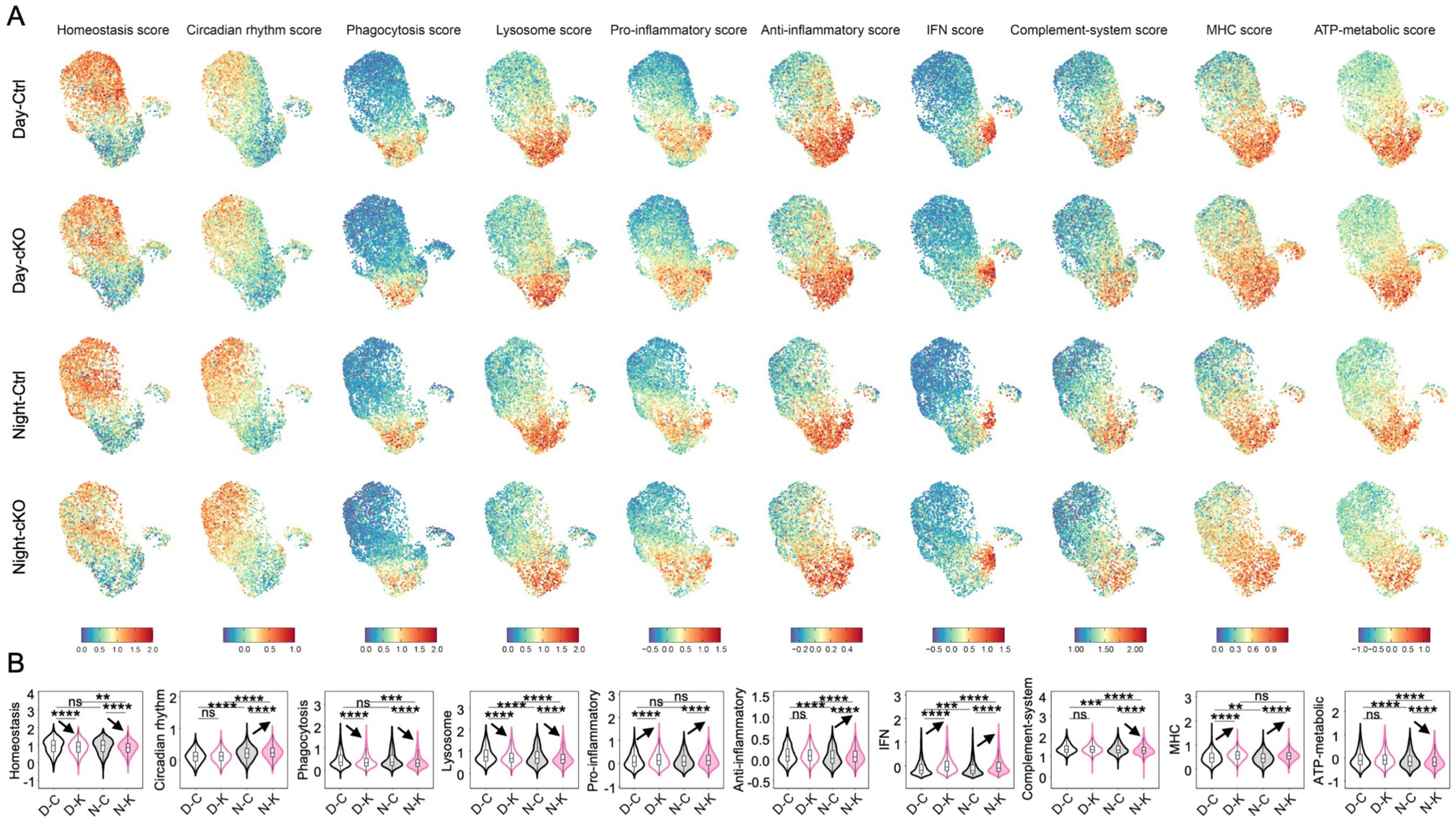
Functional profiling of microglia reveals diurnal and *Adrb2*-dependent modulation of transcriptional states. **(A)** UMAP plots visualization of functional signature scores across D-C, D-K, N-C and N-K groups. **(B)** Violin plots showing the functional signature score in microglia across the four experimental groups. Scores were calculated using AddModuleScore. **, *P*<0.01, ***, *P*<0.001, ****, *P*<0.0001, One-way ANOVA with Tukey’s HSD post-hoc test was performed.

**Figure S13.**
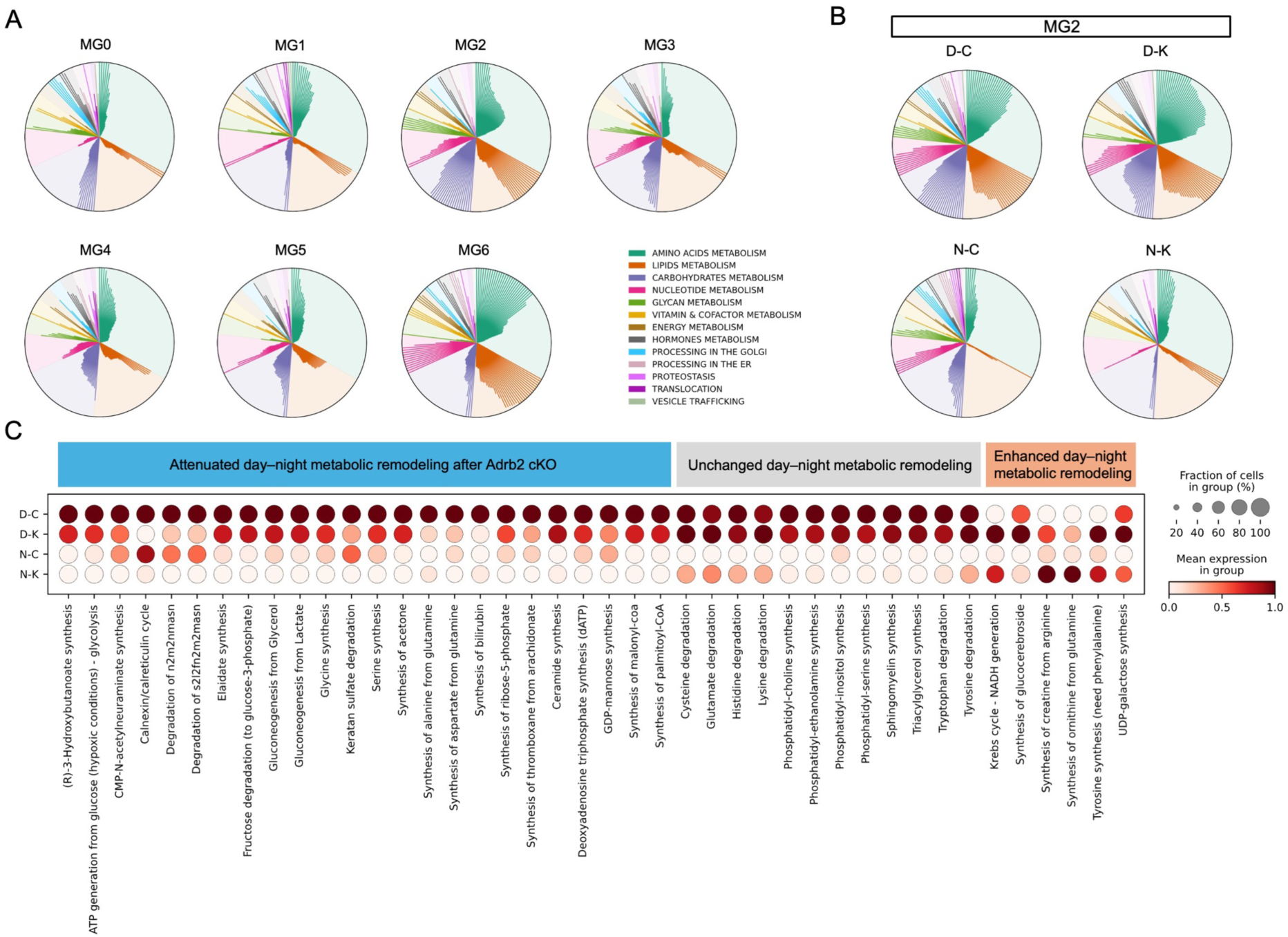
Day–night remodeling of microglial metabolic programs in control and *Adrb2* cKO mice. **(A)** Radial plots showing scCellFie-inferred metabolic task activities across microglial clusters (MG0–MG6). Metabolic tasks were grouped into functional categories, with colors indicating metabolic categories and sector length representing their relative contribution. **(B)** Radial plots showing scCellFie-inferred metabolic task activities in MG2 across D-C, D-K, N-C, and N-K groups. Metabolic tasks are summarized by functional categories as indicated. **(C)** Dot plot showing scCellFie-inferred metabolic task activities in MG2 microglia across D-C, D-K, N-C, and N-K groups. Dot size indicates the fraction of cells with activity of each metabolic task, and color intensity represents the normalized activity score.

## Movie

**Movie 1.** Longitudinal in vivo two-photon imaging of neuronal and microglial dynamics in control and cytoTDP-43 cortex, acquired every 2 days from days 16–30 after AAV injection. Left, representative footage showing the awake, head-fixed mouse during two-photon imaging, with the animal moving on the treadmill. Right, Neurons and microglia remained stable in control cortex, whereas cytoTDP-43-expressing neurons underwent progressive fluorescence accumulation, nuclear changes, somatic shrinkage and eventual signal loss followed by microglial engulfment and clearance.

**Movie 2.** Microglial depletion by PLX5622 abolishes neuronal corpse clearance. Longitudinal two-photon imaging from days 16–30 after AAV injection shows persistence of neuronal remnants in the absence of microglia, whereas PLX5622 withdrawal leads to microglial repopulation and subsequent clearance of accumulated remnants.

**Movie 3.** Multi-timescale in vivo imaging reveals the detailed dynamics of microglial phagocytosis of degenerating neurons. Imaging at progressively higher temporal resolutions, from 2-day to 3-min intervals, provides increasingly detailed visualization of microglial engulfment of degenerating neurons.

**Movie 4.** Microglial clearance varies across host rest–activity state. During the host resting/light phase, microglia engulfed more degenerating neurons and completed clearance more efficiently (white arrowheads), whereas engulfment and clearance completion were reduced during the host active/dark phase (purple arrowheads).

**Movie 5.** Transition from wakefulness to general anesthesia rapidly enhances microglial phagocytic activity.

## Table

**Table 1.** Statistical details for all data panels in the study.

**Table 2.** Differential gene expression analysis of microglia in the CBG-seq dataset.

**Table 3.** GO enrichment analysis of microglia in the CBG-seq dataset.

**Table 4.** GO enrichment analysis of microglia in scRNA-seq dataset.

**Table 5.** Marker genes used to identify major cell types in scRNA-seq dataset.

**Table 6**. Marker genes used to define microglial subclusters in scRNA-seq dataset.

**Table 7**. Differential gene expression analysis of microglia in scRNA-seq dataset.

**Table 8**. GO enrichment analysis of microglia in scRNA-seq dataset.

**Table 9.** Gene-signature scoring of microglial functional programs.

**Table 10.** Inferred metabolic-task activity profiles of microglia across experimental groups.

## Methods

### Animals and ethics statements

*Cx3cr1-GFP* (B6.129P2(Cg)-*Cx3cr1*^tm1Litt/J^, JAX:005582), *Tmem119-CreER* (BL/6.*Tmem119*^em1(cre/ERT2)Gfng/J^, JAX:031820), *Cx3cr1-CreER* (B6.129P2(Cg)-*Cx3cr1*^tm2.1(cre/ERT2)Litt/WganJ^, JAX:021160), and C57BL/6J mice (JAX:000664) were purchased from The Jackson Laboratory. *Adrb2^flox/flox^* mice were generated by GemPharmatech using CRISPR/Cas9-mediated genome editing. Briefly, sgRNAs were transcribed in vitro and co-microinjected with Cas9 mRNA and a donor vector into fertilized C57BL/6JGpt mouse zygotes. Injected embryos were transferred into pseudopregnant females to obtain F0 founder mice, which were identified by PCR and sequencing. Germline-transmitted F1 mice were generated by crossing positive F0 founders with C57BL/6JGpt mice. *Cx3cr1-GFP* mice were crossed with *Adrb2^flox/flox^* mice to generate *Adrb2^flox/flox^*::*CX3CR1^GFP/+^*mice, which were subsequently crossed with *Tmem119-CreER* mice to obtain *Adrb2^flox/flox^::Tmem119^CreER/+^:: Cx3cr1^GFP/+^*mice. In parallel, *Cx3cr1-CreER* mice were crossed with *Adrb2^flox/flox^*mice to generate *Adrb2^flox/flox^::Cx3cr1^CreER/+^ mice*. *Cx3cr1^GFP/GFP^* and *Cx3cr1^CreER/CreER^* mice were each crossed with C57BL/6J mice to generate *Cx3cr1^GFP/+^* and *Cx3cr1^CreER/+^* mice, respectively. Genotyping was performed by PCR using genomic DNA isolated from mouse tail biopsies. For *Cx3cr1-GFP* mice, the primers used were wild-type forward 5′-GTCTTCACGTTCGGTCTGGT-3′, mutant forward 5′-CTCCCCCTGAACCTGAAAC-3′, and common reverse 5′-CCCAGACACTCGTTGTCCTT-3′. For *Tmem119-CreER* mice, primers included wild-type forward 5′-CAGTATGTGGGGTCACTGAAGA-3′, mutant forward 5′-ATCGCATTCCTTGCAAAAGT-3′, and common reverse 5′-ACTTGGGGAGATGTTTCCTG-3′. For *Cx3cr1-CreER* mice, genotyping was performed using a common forward primer 5′-AAGACTCACGTGGACCTGCT-3′, together with wild-type reverse 5′-AGGATGTTGACTTCCGAGTTG-3′ and mutant reverse 5′-CGGTTATTCAACTTGCACCA-3′. *Adrb2^flox/flox^*alleles were identified using the primers forward 5′-ACTGCTCCAAGAAGCAGACTCTG-3′ and reverse 5′-GTCGTTGTCATCATCATCACTGTG-3′. All mouse lines were subsequently maintained and bred at the School of Medicine, South China University of Technology. Both male and female mice were used in this study unless otherwise specified. Mice were used at approximately 2 months of age for all experiments. Animals were housed in a specific pathogen-free (SPF) facility under controlled temperature (21 ± 1 °C) and humidity (45-60%), with ad libitum access to food and water. Mice were maintained on a 12 h light-dark cycle, with lights on from 7:00 to 19:00. All experimental procedures involving animals were approved by the Institutional Animal Care and Use Committee of the School of Medicine, South China University of Technology (approval number: AEC-2020025), and were conducted in accordance with institutional guidelines and relevant national regulations.

### Sleep deprivation and circadian inversion

Mice were maintained under a standard light-dark cycle (lights on at 07:00 and off at 19:00) unless otherwise specified. For sleep deprivation, mice were subjected to a mechanical sleep deprivation paradigm using a motorized sleep deprivation apparatus (SITRANTECH, China) consisting of a cylindrical chamber (30 cm in diameter and 45 cm in height) equipped with a motor-driven rotating interference bar (29 cm in diameter). Mice were placed in the apparatus daily during the light phase (10:00-16:00), during which the interference bar rotated at 2 rpm to maintain wakefulness. Food and water were available ad libitum throughout the procedure. Animals were continuously monitored to ensure sustained wakefulness. Sleep deprivation was performed for 7 consecutive days before experimental procedures. For circadian inversion, mice at approximately one month of age were transferred to a reversed light-dark cycle, in which the dark phase and light phase were inverted (lights off at 07:00 and on at 19:00). Mice were maintained under the inverted cycle for at least 4 weeks to allow circadian entrainment before subsequent experiments.

### Drug administration

For pharmacological manipulation of β2-adrenergic signaling, mice received intraperitoneal (i.p.) injections of the ADRB2 antagonist Zenidolol hydrochloride ICI-118,551 (MedChemExpress, HY-13951; 10 mg kg⁻¹), or the ADRB2 agonist formoterol fumarate (MedChemExpress, HY-B0010; 2 mg kg⁻¹). Both compounds were dissolved in 5% DMSO and 95% saline. Selective ablation of locus coeruleus (LC) noradrenergic neurons was achieved by i.p. administration of the neurotoxin DSP-4 hydrochloride (MedChemExpress, HY-103210; 50 mg kg⁻¹) dissolved in saline. For chemogenetic manipulation, clozapine N-oxide (CNO; MedChemExpress, HY-17366; 1 mg kg⁻¹) was administered i.p. in 5% DMSO and 95% saline. To induce Cre recombinase activity, tamoxifen (MedChemExpress, HY-13757A) was dissolved in corn oil at a concentration of 20 mg mL⁻¹ and administered i.p. at 75 mg kg⁻¹. For pharmacological depletion of microglia, mice were fed a diet containing the CSF1R inhibitor PLX5622 (1200 ppm; SYSE), while control mice received an AIN-76A control diet (SYSE). The PLX5622 diet was custom formulated by incorporating 1.2 g PLX5622 per kg of AIN-76A chow, and mice had ad libitum access to food.

### AAV injection and cranial window surgery

Age-matched mice (∼2 months old) were used for all surgical procedures. Mice were deeply anesthetized in an induction chamber with 5% isoflurane until respiration slowed and no response to paw pinch was observed and were subsequently secured in a stereotaxic frame (RWD, 68804). Anesthesia was maintained throughout surgery with 1-1.2% isoflurane delivered at an airflow rate of 0.5 L min^-1^. Body temperature was maintained using a thermostatically controlled heating pad. After removal of scalp hair and disinfection with povidone-iodine, the scalp was incised along the midline. The skull surface was cleaned using 3% hydrogen peroxide and rinsed with sterile saline. The skull was leveled such that the height differences between bregma and lambda, as well as between the left and right hemispheres, did not exceed 0.03 mm. Using bregma as the reference point, a small craniotomy was made above the primary motor cortex (M1; AP +1.5 mm, ML ±1.8 mm). Viral vectors (300 nL) were delivered at a rate of 60 nL min^-1^ using a microsyringe pump, with the injection needle positioned at DV −1.0 mm. Following injection, the needle was left in place for an additional 10 min before slow withdrawal. The scalp was sutured with nylon thread, and mice were returned to their home cages after recovery from anesthesia. For cranial window implantation, the scalp was completely removed to expose the skull, and connective tissue was thoroughly cleared. After rinsing with sterile saline, a circular craniotomy (∼3.5 mm diameter) was thinned and opened above the right primary somatosensory cortex (S1). The exposed cortical surface was continuously irrigated with sterile saline and covered with saline-soaked gelatin sponge to maintain moisture. Viral injections were performed at S1 (AP −1.5 mm, ML −1.8 mm, DV −0.75 mm). After injection, the window was cleaned and temporarily covered with gelatin sponge. The skull surface was dried and coated with adhesive (3M ESPE Single Bond Universal, 3M, 41282), which was rapidly cured under blue light. A 4-mm circular glass coverslip (thickness ∼0.17 mm) was placed over the craniotomy and sealed with flowable resin (3M, Z350 XT) under blue light illumination. For awake two-photon imaging, a custom-designed metal head ring (SITRANTECH, China) was affixed to the skull using dental resin, ensuring that the cranial window was centered and aligned with the imaging axis. After surgery, isoflurane was discontinued, and mice were returned to their home cages once fully ambulatory. To manipulate locus coeruleus (LC) activity, chemogenetic approaches were employed. AAV vectors encoding inhibitory hM4Di DREADDs were stereotaxically injected (200 nL) into the LC (AP −5.4 mm, ML ±0.9 mm, DV −3.5 mm) to selectively inhibit LC neurons.

The following adeno-associated viruses (AAVs) were used in this study:

AAV9[ssAAV.CaMKIIα.-mTDP-43(mutNLS)-linker-mScarlet.WPRE.SV40pA] (PackGene; Custom-made; 1 × 10¹² GC mL^-1^);
AAV9[ssAAV.CaMKIIα-mScarlet.WPRE.SV40pA] (PackGene; Custom-made; 1 × 10¹² GC mL^-1^);
AAV9[PFD-rAAV-hSyn-jGCaMP7f-WPRE-hGH polyA] (BrainVTA, Cat# PT-1887; 2 × 10¹² VG mL^-1^);
AAV9[PFD-rAAV-TH-Cre-WPRE-hGH polyA] (BrainVTA, Cat# PT-3624; 5 × 10¹⁰ VG mL^-1^); AAV9[rAAV-EF1α-DIO-hM4D(Gi)-mCherry] (Braincase, Cat# BC-0155; 1 × 10¹² VG mL^-1^);
AAV9[rAAV-hSyn-NE2h] (Braincase, Cat# BC-0268; 2 × 10¹² VG mL^-1^).

Following surgery, mice were placed on a 37 °C heating pad until recovery and then returned to their home cages. Mice were allowed at least 2 weeks for viral expression and were habituated to head fixation on the treadmill platform for 3 consecutive days (30 min day⁻¹) before imaging.

### Tissue processing and cryo-sectioning

Mice were deeply anesthetized in an induction chamber with 5% isoflurane and secured before transcardial perfusion. Animals were first perfused with 0.01 M phosphate-buffered saline (PBS) to remove blood, followed by perfusion with 4% paraformaldehyde (PFA; Biosharp, BLS39A) prepared in 0.01 M PBS for pre-fixation. Brains were carefully dissected and post-fixed by immersion in 4% PFA at 4 °C overnight. Tissues were then subjected to sucrose dehydration by sequential incubation in 20% and 30% sucrose solutions (both prepared in 0.01 M PBS) for 24 h at each concentration at 4 °C. After completion of sucrose dehydration, brains were embedded in optimal cutting temperature compound (OCT, RWD) and frozen. The embedded tissues were equilibrated in a cryostat (Leica, CM1950) at −25 °C prior to sectioning. Coronal sections were cut at a thickness of 30 μm and collected into 24-well plates containing 0.01 M PBS for subsequent staining.

### Immunostaining

Free-floating tissue sections were used for both immunofluorescence and immunohistochemistry. Sections were rinsed in 0.01 M Tris-buffered saline (TBS) for 10 min and blocked in blocking solution consisting of 5% goat serum (Solarbio, SL038) and 0.5% Triton X-100 (Sigma-Aldrich, T8787) in 0.01 M TBS for 1 h at room temperature. After removal of the blocking solution, sections were incubated with primary antibodies diluted in antibody dilution buffer (0.5% TBST; 0.01 M TBS containing 0.5% Triton X-100) at 4 °C overnight. Sections were then brought to room temperature and washed three times with 0.01 M TBS for 10 min each. For immunofluorescence, sections were incubated with appropriate fluorophore-conjugated secondary antibodies diluted in 0.5% TBST for 2 h at room temperature. After three washes in 0.01 M TBS (10 min each), sections were counterstained with 4′,6-diamidino-2-phenylindole (DAPI; Beyotime, C1006) for 15 min, followed by three additional washes. Sections were then mounted onto glass slides, air-dried at room temperature, and coverslipped with an antifade mounting medium (Sigma-Aldrich, F4680) for subsequent imaging. For immunohistochemistry, sections were pretreated with 3% hydrogen peroxide for 15 min prior to blocking to quench endogenous peroxidase activity, followed by rinsing in 0.01 M TBS for 10 min. Sections were then processed using the same blocking and primary antibody incubation procedures as described above. An HRP-conjugated secondary antibody detection system was used, and immunoreactivity was visualized using 3,3′-diaminobenzidine (DAB) as the chromogenic substrate. Primary antibodies used in this study included rabbit anti-TDP-43 (10782-2-AP, 1:500; Proteintech), guinea pig anti-NeuN (ABN90, 1:1500; Sigma-Aldrich), rabbit anti-IBA1 (ab178846, 1:1000; Abcam), rat anti-IBA1 (ab283346, 1:2000; Abcam), rat anti-CD68 (ab53444, 1:500; Abcam) and rabbit anti-P2Y12 (AS-55043A, 1:800; ANASPEC). For immunofluorescence, the following fluorophore-conjugated secondary antibodies were used: goat anti-rabbit IgG (H+L) Alexa Fluor 488 (A-11008, 1:800; Thermo Fisher Scientific), goat anti-guinea pig IgG (H+L) Alexa Fluor 647 (106-605-003, 1:500; Jackson ImmunoResearch), goat anti-rat IgG (H+L) Alexa Fluor 488 (A11006, 1:500; Thermo Fisher Scientific), and goat anti-rat IgG (H+L) Alexa Fluor 647 (A21247, 1:500; Thermo Fisher Scientific). For immunohistochemistry, an HRP-conjugated goat anti-rabbit IgG (H+L) secondary antibody (SA00001-2, 1:500; Proteintech) was used.

### Fluorescence in situ hybridization

Combined immunofluorescence and RNA fluorescence in situ hybridization (RNA-FISH) was used to evaluate the efficiency of *Adrb2* deletion in microglia. Mouse brains were fixed in 4% PFA and cryoprotected in a graded sucrose series. Coronal sections (10 μm) were prepared using a cryostat and mounted onto positively charged glass slides. Sections from knockout and control mice were mounted on the same slide to minimize technical variability. Slides were dried at 37 °C for 1 h and processed for immunofluorescence staining to label microglia as described above. To preserve RNA integrity, RNase inhibitor (1 U μl^-1^; Takara, cat# 2313A) was included in both the blocking and primary antibody incubation buffers. Immediately after immunofluorescence staining, RNA in situ hybridization was performed using the PinpoRNA™ 2.0 Tissue RNA ISH Detection Kit (Pinpoease, PIF2000) according to the manufacturer’s instructions. Sections were sequentially subjected to protease digestion, probe hybridization, signal amplification, and fluorescent labeling to enable detection and quantification of target transcripts. The mouse *Adrb2* probe was designed based on the NCBI reference sequence (Gene ID: 11555; RefSeq NM_007420.3).

### Image acquisition

Histological images were acquired using a laser scanning confocal microscope (LSM 900, Zeiss), with either single-plane (XY) or z-stack (XYZ) acquisition at 10× or 40× objectives. Whole-slide imaging was performed using a slide scanner (Vectra Polaris, Akoya Biosciences) at ×20 magnification.

### Transmission electron microscopy

Mice were deeply anesthetized with isoflurane (5% for induction) and transcardially perfused with ice-cold saline to remove blood, followed by fixation with a precooled solution containing 2% paraformaldehyde (Sigma-Aldrich, 16005) and 2.5% glutaraldehyde (Sigma-Aldrich, 340855) diluted in 0.01 M PBS. Brains were rapidly dissected, and regions of interest were cut into ∼1 mm³ blocks and post-fixed overnight at 4 °C in 2.5% glutaraldehyde prepared in 0.01 M PBS. Samples were rinsed several times in 0.01 M PBS, post-fixed with 1% osmium tetroxide for 2 h, dehydrated through a graded ethanol series, and embedded in epoxy resin. Ultrathin sections (70-90 nm) were prepared using an ultramicrotome, collected on copper grids, and sequentially stained with uranyl acetate and lead citrate (5 min each). Sections were examined and imaged using a transmission electron microscope (HT7800, Hitachi). Sample post-processing and imaging were performed at the Pearl River Fisheries Research Institute.

### Two photon imaging

Beginning 14 days after cranial window implantation, mice were habituated to head fixation on a rotating disk treadmill (SITRANTECH, China) while allowing unrestricted movement of the limbs (30 min daily for 7 consecutive days). Imaging was performed on a Leica STELLARIS 8 two-photon microscope equipped with a 25× water-immersion objective (NA 0.95). Image stacks (1024 × 1024 pixels; 221 × 221 × 100 μm; 101 optical sections at 1 μm intervals) were acquired 2-day intervals from 16 to 30 days after viral injection to monitor microglial and neuronal dynamics. GFP and mScarlet were excited at 924 nm and 1040 nm, respectively. Fluorescence emission was collected using a combination of variable dichroics and band-pass filters at 480-530 nm (GFP) and 580-650 nm (mScarlet). For high-temporal-resolution imaging, time-lapse recordings were acquired at intervals ranging from 3 min to 2 h, with continuous imaging sessions lasting 2–30 h depending on the experimental design. For longitudinal imaging performed at 2-day, 12-h, or 2-h intervals, mice were returned to their home cages between imaging sessions, and identical imaging fields were relocated in subsequent sessions using surface vascular landmarks together with neuronal morphology. For short-interval time-lapse imaging (3 or 20 min), mice remained under the microscope throughout the continuous recording session.

### Image analysis

Histological and two-photon images were analyzed using Fiji (ImageJ, NIH) unless otherwise indicated. For histological analyses, confocal z-stacks acquired with a 40× objective were projected as maximum-intensity projections over a depth of 20 μm. Cell density was quantified from projected images using the Analyze Particles function with identical thresholding parameters applied across all samples. For two-photon image stacks, motion artifacts induced by mouse movement were corrected using the StackReg plugin. Multichannel fluorescence images were first separated into individual channels using the Split Channels function. Cell segmentation was performed on the channel defining cell boundaries. Images were converted to grayscale and thresholded using the Otsu method (Image/Adjust /Threshold) to generate binary masks. Because the mScarlet signal was predominantly localized to the plasma membrane, binary images were further processed with morphological operations to improve segmentation accuracy. Internal holes were filled using the Fill Holes function, and residual noise or small artifacts were removed using Open and Close operations (Process/Binary). Cell size was quantified using the Analyze Particles function (Analyze/Analyze Particles). Three-dimensional reconstructions of microglia-neuron interactions during phagocytosis were generated from two-photon image stacks using the Surface function in Imaris (Bitplane). Image analyses were conducted blindly by two independent investigators to reduce bias.

### Quantification of microglial phagocytosis

**The microglial engulfment index** was defined as the number of newly initiated microglial engulfment events occurring between two consecutive imaging sessions within a given field of view. Values were normalized to the first imaging session of each longitudinal recording to facilitate comparisons across day and night. **Microglial phagocytic index** was quantified as the fraction of the neuronal area occupied by GFP⁺ microglial signals relative to the mScarlet⁺ neuronal area. This metric provides a continuous estimate of phagocytic progression, ranging from approximately 0 before microglial enclosure of the neuron to approximately 1 after complete engulfment and disappearance of the neuronal mScarlet signal. This index reflects the extent of phagocytosis achieved by an individual microglial cell during neuronal corpse clearance. **Cumulative phagocytic index** was calculated by summing the phagocytic indices of all individual microglia within a given imaging field. This metric reflects the overall phagocytic activity of the local microglial population and represents the total extent of neuronal engulfment within the field of view.

### Behavioral classification of microglial phagocytosis

Microglial phagocytic behavior was classified into five sequential stages based on longitudinal in vivo two-photon imaging. Stage 0 (Prepare) was defined as the initial contact of a microglial process with a degenerating neuron without evident morphological polarization. Stage 1 (Reach) was defined by directed extension of microglial processes toward the neuronal soma. Stage 2 (Approach) was defined as translocation of the microglial soma toward the target neuron while maintaining process contact. Stage 3 (Isolate) was defined by complete encapsulation of the neuronal soma by the microglial cell body and processes, resulting in physical isolation of the neuron from the surrounding parenchyma. Stage 4 (Digest) was defined as progressive degradation and clearance of the neuronal corpse following engulfment, accompanied by the disappearance of mScarlet-positive neuronal material.

### Neuronal calcium Signal Analysis

Time-series images of neuronal calcium activity were motion-corrected using the StackReg plugin in ImageJ (Fiji). An average-intensity projection of the time series was generated to facilitate region-of-interest (ROI) selection. ROIs were manually drawn over neuronal somata using the circular selection tool, restricted to neurons that were positive for both GCaMP and mScarlet signals. Mean fluorescence intensity values (F) for each ROI were extracted using the Time Series Analyzer plugin. To minimize the influence of slow fluorescence drift and transient calcium spikes on baseline estimation, baseline fluorescence (F₀) was defined as the 25th percentile of fluorescence values across the recording. Fluorescence traces were then converted to normalized calcium signals (ΔF/F₀) according to:

Δ F/F_0_ = (F - F_0_)/F_0_

Calcium events were defined as fluorescence transients in which ΔF/F₀ exceeded two standard deviations above baseline for at least two consecutive frames. Event frequency was calculated as the number of calcium events per minute.

### Behavior-guided SMART-seq

#### Slice preparation

CX3CR1^GFP/+^ mice were subjected to pathological modeling in the primary motor cortex (M1). 25 days after model induction, animals were used for microglial transcriptomic analysis. Mice were deeply anesthetized with 5% isoflurane and transcardially perfused with ice-cold, oxygenated cutting buffer to remove circulating blood cells. Brains were rapidly dissected and immediately transferred into ice-cold, oxygenated cutting buffer containing (in mM): 120 choline chloride, 2.5 KCl, 1.25 NaH₂PO₄, 7 MgCl₂, 0.5 CaCl₂, 25 NaHCO₃ and 10 D-glucose. Coronal brain slices (300 μm thickness) encompassing the modeled M1 region were prepared using a vibratome (Leica, VT1200S). All slices were briefly equilibrated in oxygenated recording buffer containing (in mM): 126 NaCl, 3 KCl, 1.25 NaH₂PO₄, 1 MgSO₄, 2 CaCl₂, 26 NaHCO₃ and 10 D-glucose, and subsequently incubated in oxygenated recording buffer at 37.0 ± 0.5 °C for 0.5-1 h prior to downstream processing. **Single-cell isolation**. To couple microglial behavior with transcriptomic identity, individual microglia were visually identified in acute brain slices based on their morphological and phagocytic states and directly aspirated into patch pipettes for downstream SMART-seq analysis. All work surfaces were thoroughly cleaned with DNA-OFF (Takara, #9036) and RNase Zap (ThermoFisher Scientific, AM9780) prior to experiments. Patch-clamp pipettes (outer diameter, 1.5 mm; inner diameter, 0.86 mm) were freshly pulled on the day of use. After incubation, brain slices were transferred to the stage of a whole-cell patch-clamp microscope and stabilized using a fine nylon mesh mounted on a platinum ring. Throughout the procedure, slices were continuously superfused with oxygenated recording buffer at a constant flow rate to maintain tissue viability, and the total handling time for each slice did not exceed 1 h. The pipette tip was filled with 0.5 μl RNase-free intracellular solution containing (in mM): 123 potassium gluconate, 12 KCl, 10 HEPES, 0.2 EGTA, 4 MgATP, 0.3 NaGTP and 10 sodium phosphocreatine, supplemented with 20 μg ml^-1^ glycogen and 1 U ul^-1^ recombinant RNase inhibitor. Target microglial cells were visually identified based on morphological criteria using a micromanipulator. Positive pressure was applied to gently displace the surrounding extracellular matrix, followed by negative pressure to aspirate a single cell into the pipette. Immediately after cell capture, the pipette containing the single cell was withdrawn and its contents were expelled into a PCR tube containing pre-chilled CDS Sorting Solution (total volume, 11 μl; 0.95 μl 10× lysis buffer, 0.05 μl RNase inhibitor and 10 μl nuclease-free water). Care was taken to avoid bubble formation to minimize cell rupture and RNA degradation. Samples were snap-frozen in liquid nitrogen and stored until subsequent cDNA amplification and library preparation. **cDNA synthesis and amplification**. To generate high-quality, high-yield cDNA, single-cell reverse transcription and cDNA amplification were performed using the SMART-Seq HT Kit (Takara, Cat. #634438) according to the manufacturer’s instructions with minor modifications. All work surfaces were decontaminated with DNA-OFF and RNase Zap prior to use, and all procedures were carried out in a fume hood under continuous ventilation to minimize nucleic acid contamination. PCR tubes containing liquid nitrogen-frozen samples were thawed on ice. Subsequently, 1 μl of 3′ SMART-Seq CDS Primer II A was added to each tube, followed by gentle mixing and brief centrifugation. Samples were incubated at 72 °C for 3 min in a preheated thermal cycler and immediately placed on ice for 2 min. A total of 12.5 μl One-Step Master Mix (0.7 μl nuclease-free water, 8 μl One-Step Buffer, 1 μl SMART-Seq HT Oligonucleotide, 0.5 μl RNase inhibitor, 0.3 μl SeqAmp DNA polymerase and 2 μl SMARTScribe reverse transcriptase) was then added to each reaction. After gentle mixing and centrifugation, reverse transcription and cDNA amplification were performed in a thermal cycler. Amplified cDNA was purified using AMPure XP beads (Beckman Coulter Life Sciences, #A63880), yielding a final volume of 17 μl purified cDNA. cDNA concentration was quantified using the Equalbit 1× dsDNA HS Assay Kit (Vazyme, #EQ121) on a Qubit 4 fluorometer (Thermo Fisher Scientific). Fragment size distribution and cDNA quality were assessed using the High Sensitivity DNA Kit (Agilent, #5067-4626) on an Agilent 2100 Bioanalyzer (Agilent). Only samples passing quality control were used for subsequent library preparation and sequencing. **Library construction and sequencing**. For Illumina high-throughput sequencing, sequencing libraries were prepared using the TruePrep DNA Library Prep Kit V2 for Illumina (Vazyme, TD503) in combination with the TruePrep Index Kit V2 for Illumina (Vazyme, TD202) to generate dual-indexed paired-end libraries. Library purification and size selection were performed using AMPure XP beads (Beckman Coulter Life Sciences, #A63880) according to the manufacturer’s instructions. Constructed libraries were subjected to quality control by Novogene prior to sequencing. Libraries that passed quality assessment were sequenced on an Illumina NovaSeq X Plus platform with paired-end 150-bp reads (PE150), generating approximately 2 Gb of data.

### Single-cell RNA sequencing

#### Brain tissue preparations

Two independent batches of brain tissues were processed for single-cell RNA sequencing in this study. In the first batch, wild-type C57BL/6 mice received stereotaxic injections of either CamkIIα-TDP-43-mScarlet or control CamkIIα-mScarlet virus into the primary motor cortex (M1; anteroposterior (AP), +1.5 mm; mediolateral (ML), ±1.8 mm; dorsoventral (DV), −1.0 mm). In the second batch, Adrb2^f/f^::CX3CR1^CreER/+^ (Adrb2 cKO) mice and their littermate controls (CX3CR1^CreER/+^) were injected with CamkIIα-TDP-43-mScarlet virus into the M1 region. Brain tissues were collected 25 days after viral injection. To investigate circadian-associated transcriptional changes in microglia, tissues from the first batch were collected during the daytime (11:00-12:00), whereas tissues from the second batch were collected both during the daytime (11:00-12:00) and nighttime (23:00-24:00). To preserve tissue freshness, mice were deeply anesthetized with 5% isoflurane and transcardially perfused with ice-cold, oxygenated cutting buffer. After complete clearance of blood, brains were rapidly dissected, and the M1 region was isolated using a double-edged razor blade. Tissue samples were temporarily stored in pre-chilled MACS Tissue Storage Solution (Miltenyi Biotec, 130-100-008) prior to further processing. For single-cell suspension preparation, each experimental group consisted of three biological replicates, with each replicate pooled from the bilateral M1 regions of approximately three mice. **Single-cell suspension preparation.** Freshly dissected M1 brain tissues were processed for single-cell suspension preparation. After removal of meninges and connective tissues, samples were rinsed with cold PBS and minced into ∼1-2 mm³ fragments in serum-free DMEM. Tissue fragments were enzymatically digested in DMEM containing papain and DNase I at 37 °C for approximately 20 min with gentle agitation. Enzymatic digestion was terminated by adding DMEM supplemented with 10% fetal bovine serum, followed by filtration through a 70-μm cell strainer to remove undigested debris. Cell suspensions were centrifuged at 350 g for 5 min at 4 °C, resuspended, and subjected to red blood cell lysis using eBioscience™ 1× Red Blood Cell Lysis Buffer (ThermoFisher, 00-4333-57) according to the manufacturer’s instructions. After washing, cells were gently resuspended in DMEM containing 0.04% BSA and 0.05 mM EDTA. Myelin debris was subsequently removed using Debris Removal Solution (Miltenyi Biotec, 130-109-398) according to the manufacturer’s instructions. Cells were then washed, centrifuged, and filtered through a 40-μm cell strainer. Final cell suspensions were resuspended in DMEM containing 0.04% BSA and 0.05 mM EDTA, and cell concentration and viability were assessed using trypan blue staining. Only samples with viability greater than 80% and a concentration of 1,000-2,000 cells/μl were used for downstream single-cell RNA sequencing. **10x Genomics and sequencing.** After confirming that the quality of the single-cell suspension met the required criteria, single-cell capture and library preparation were performed using the Chromium Next GEM Single Cell 3′ GEM, Library & Gel Bead Kit v3.1 together with the Chromium Next GEM Chip G Single Cell Kit (10x Genomics), following the manufacturer’s instructions. Briefly, single-cell suspensions were counted and assessed for viability, and approximately 13,000-15,000 cells per sample were loaded onto the Chromium Controller to generate gel bead-in-emulsions (GEMs), in which individual cells were co-encapsulated with barcoded gel beads carrying unique molecular identifiers (UMIs). Reverse transcription was carried out within the GEMs, after which emulsions were broken and cDNA was recovered, amplified, and used for 3′ gene expression library construction according to the standard protocol. Final libraries were quality-controlled and quantified prior to sequencing. Libraries were sequenced on an Illumina NovaSeq X Plus platform in paired-end mode, achieving a sequencing depth of approximately 100 Gb per sample. Single-cell suspension preparation and 10x Genomics library construction and sequencing were performed by GENE DENOVO Biotechnology.

### Analysis of smart-seq data

Raw FASTQ files from -SMART-seq were processed for quality control and adapter trimming using Trim Galore (v0.6.10) with default parameters. Cleaned reads were aligned to the mouse reference genome GRCm39 using the splice-aware aligner HISAT2 (v2.2.1). The resulting Sequence Alignment/Map (SAM) files were converted to Binary Alignment/Map (BAM) format, sorted by genomic coordinates, and indexed using SAMtools (v1.17). Gene-level read counts were generated with HTSeq-count (v2.0.2) in ‘union’ mode against the Mus_musculus.GRCm39.113.gtf annotation file, yielding a raw count matrix for downstream analysis.

All subsequent analyses were performed in R (v4.5.1). Low-abundance genes (total counts < 10 across all samples) were filtered out. Differential expression analysis was conducted using DESeq2 (v1.48.1), with multiple testing correction applied via the Benjamini-Hochberg (BH) method. Data quality was assessed by principal component analysis (PCA) and a sample correlation heatmap, both generated from variance-stabilizing transformed (VST) data; the heatmap visualized the top 100 most highly expressed genes using the pheatmap package (v1.0.13).

Genes with an adjusted p-value < 0.05 and a |log2FC| (FC: Fold change) > 0.5 were considered significantly differentially expressed. Gene Ontology (GO) enrichment analysis for upregulated and downregulated DEGs was performed separately using clusterProfiler (v4.16.0) and the org.Mm.eg.db (v3.21.0) database, with BH adjustment. Eight microglial functional signatures were defined based on published marker genes and functional annotations. Gene expression within each signature was visualized using VST-normalized expression data and compared by one-way ANOVA with Tukey’s HSD post-hoc test. Finally,a weighted gene co-expression network analysis (WGCNA) was applied to the top 5,000 most highly expressed genes using the WGCNA package (v1.73). A soft threshold power was selected to achieve a scale-free topology fit (R² ≥ 0.9) for an unsigned-hybrid network. GO enrichment analysis was performed for each module, and the top 25 genes (ranked by module membership) were used for network visualization with ggplot2 (v4.0.1).

### Analysis of scRNA-seq

For 3’ single-cell RNA-seq data, sample demultiplexing, barcode processing, and gene counting were performed using the Cell Ranger 10x Genomics (v8.0.0). Briefly, sample-specific FASTQ files were generated by demultiplexing 8 bp index reads. Reads were aligned to the GRCm39 genome using the STAR aligner. After alignment, barcodes and UMIs were filtered and corrected, and a gene-barcode matrix was generated, retaining only confidently mapped, non-PCR duplicate reads with valid barcodes and UMIs.

Downstream analyses were performed using Seurat (v5.3.1). Cells were filtered based on quality control metrics (gene number <200 or >6000, UMI counts <500 or >20,000, and mitochondrial RNA proportion >10%). Ambient RNA contamination and doublets were removed using decontX (v1.6.0; contamination score >0.8) and DoubletFinder (v2.0.6; pN = 0.25, optimal pK), respectively. Filtered datasets were normalized using SCTransform and subjected to PCA, clustering, and UMAP visualization based on the top 30 principal components. Batch correction was performed using Harmony (v1.2.4). Cluster marker genes were identified using FindAllMarkers (Wilcoxon rank-sum test; min.pct = 0.25, logFC threshold = 0.25, only.pos = TRUE) after SCT-based preparation with PrepSCTFindMarkers. Cell types were annotated based on canonical marker genes, and clusters with ambiguous microglial and non-microglial lineage signatures were excluded.

Microglial Seurat was extracted from scRNA-seq dataset for SMART-seq transcriptome projection. Microglial cells were reprocessed using SCTransform, and contaminating endothelial cells were removed based on UCell endothelial signature scores. SMART-seq profiles were mapped onto the reference dataset using Seurat anchor-based transfer methods (FindTransferAnchors, TransferData, and IntegrateEmbeddings), followed by joint visualization of projected embeddings and cluster identities.

For sample-level comparisons, **pseudobulk differential expression analysis** was performed using DESeq2 (v1.48.1). Raw counts from the RNA assay were aggregated by biological samples using Seurat AggregateExpression, followed by filtering genes with fewer than 10 total counts. Differentially expressed genes were defined as adjusted P value < 0.05 and |log2FC| > 0.25. Heatmaps of differentially expressed genes were generated using clusterGVis (v0.99.1). **Differential gene expression analysis** was performed using Seurat FindMarkers with the MAST test (min.pct = 0.1, logfc.threshold = 0.25) for comparisons between MG1 and MG0. Genes with adjusted P value < 0.05 and |log2FC| thresholds of 0.5 were considered significant according to the corresponding comparisons.

**Gene Ontology (GO) enrichment analysis** for upregulated and downregulated DEGs was performed separately using clusterProfiler (v4.16.0) and the org.Mm.eg.db (v3.21.0) database, with BH adjustment.

**Pseudotime analysis** was performed using Monocle3 (v1.4.26). Microglial Seurat objects were converted into Monocle3 cell_data_set objects, retaining the original UMAP embeddings and cluster annotations. Trajectories were reconstructed using learn_graph with partition information disabled (use_partition = FALSE), and cluster 0 representing homeostatic microglia was defined as the root population for pseudotime ordering. Gene expression dynamics of representative microglial programs, including homeostasis, phagocytosis, lysosomal, and ATP metabolic genes, were evaluated along pseudotime using LOESS smoothing. Pseudotime distributions across experimental groups were visualized by kernel density estimation, and differences between groups were assessed using two-sided Wilcoxon rank-sum tests.

**RNA velocity** analysis was performed using Velocyto and scVelo (v0.3.3) based on spliced and unspliced transcript counts in Python (v3.13.5). Velocity dynamics were inferred using scvelo.tl.velocity after standard preprocessing.

**Differential abundance of cellular neighborhoods** was tested using MiloR (v2.4.1) with parameters: prop=0.2, k=30, d=12. Differences in abundance were tested between daytime and nighttime TDP43 conditions.

**Signature scores** were calculated using AddModuleScore, and differences were compared using a two-tailed Student’s t-test or one-way ANOVA with Tukey’s HSD correction.

**Metabolic task activities** were inferred using the scCellFie package (v0.6.0) in Python (v3.13.5) with normalized scRNA-seq expression matrices as input. Metabolic task activity profiles were compared across microglial clusters and experimental conditions. For MG2 cells, differential metabolic task activities between groups were evaluated, and activity scores were visualized using dot plots after task-wise min–max scaling.

### Statistics

Statistical analyses were carried out using GraphPad Prism (v10.1.2). Comparisons between two groups were made using unpaired two-tailed welch t-tests. Comparisons between two groups of day-night difference were made using paired two-tailed Student’s t-tests. For analyses involving three or more groups, one-way or two-way ANOVA followed by Šidák’s, Dunnett’s or Tukey’s post hoc tests was applied, as appropriate. Error bars are the mean ± s.e.m. unless stated otherwise. The exact n (number of mice or neurons) and statistical tests used are provided in the Figure legends. Statistical significance was defined as *P* < 0.05 (* or #), *P* < 0.01 (** or ##), *P* < 0.001 (*** or ###), and *P* < 0.0001 (****). For blinded quantifications, rater values were averaged per field of view and then collapsed to animal-level means to avoid pseudoreplication; these animal means were used for statistical inference.

